# Drying kinetics govern transcriptional and post-transcriptional reprogramming during seed maturation

**DOI:** 10.64898/2026.04.28.721260

**Authors:** Asif Ahmed Sami, Leo A. J. Willems, Lukman Abdulroheem, Marie-Christine Carpentier, Rémy Merret, Leónie Bentsink, Mariana A. S. Artur

## Abstract

Desiccation tolerance (DT) serves as a cornerstone for seed survival and for long-term persistence in the natural environment. DT is acquired during seed development, as seeds undergo a drastic change in internal water content during maturation drying. Although the physiological effects of drying on the acquisition of DT and other seed traits have been described, the molecular mechanisms underlying these effects have not yet been fully understood. Here, we addressed this gap by submitting maturing seeds of *Arabidopsis thaliana* L. to three different drying regimes - fast drying (FD), slow drying (SD), and a combination of both (SDFD) and studying physiological, transcriptional, and post-transcriptional responses. We found that SD not only accelerated DT acquisition but also seed maturation. Each drying regime showed a distinct transcriptional signature, with SD and SDFD showing greater global gene downregulation compared to FD. This downregulation appeared to be crucial for establishing DT in developing seeds. Interestingly, FD triggered a specific defense-related transcriptional response that was detrimental to seed longevity. Using an abscisic acid deficient mutant, we found that most of the drying-mediated transcriptional changes were largely independent of the wild-type ABA levels. On a post-transcriptional level, SD led to a major turnover of mRNA populations undergoing co-translational mRNA decay (CTRD) and promoted CTRD of stress-related genes. Overall, our study provides fundamental insights into the mechanisms by which seeds perceive and respond to drying, advancing our basic understanding of the molecular regulation of DT and seed maturation.

**Significance Statement:** Seed maturation is a critical phase of the plant life cycle when seeds acquire desiccation tolerance (DT) required for long-term storage. Drying rate, together with abscisic acid (ABA), has been implicated in this process, but whether seed development actively responds to different drying rates and how such responses are regulated has remained unclear. Here, we show that maturing seeds sense and respond to different drying regimes through distinct molecular programs, with slow drying triggering coordinated transcriptional and post-transcriptional reprogramming associated with enhanced DT. This response occurs partly independent of wild-type ABA levels, revealing drying rate as a developmental signal acting alongside hormonal regulation to direct seed maturation. These findings provide a framework for improving drying strategies and identifying molecular markers of seed quality.

## Introduction

Seeds are a vital cornerstone of modern agriculture and global food security. Seed-related traits have been the constant focal points of domestication throughout the history of major angiosperm crop species (Diamond, 2002; Meyer & Purugganan, 2013). A large share of this success can be attributed to the ability of most angiosperm seeds (called orthodox seeds) to be stored under dry conditions for prolonged periods. Although much remains unknown, seed storability, or longevity, is based on a unique biological trait called desiccation tolerance (DT). Molecular mechanisms of DT enable seeds to lose up to 90% of their water content without dying. Despite being an ancestral trait, DT is mainly restricted to reproductive structures such as spores, orthodox seeds, and pollens and vegetative tissues of resurrection plants (Hoekstra *et al*., 2001; Oliver *et al*., 2020; Marks *et al*., 2025).

In orthodox seeds, DT is acquired during the maturation phase of seed development, along with other quality traits such as germinability, dormancy, vigor and longevity (Leprince *et al*., 2016; Artur *et al*., 2024). Following embryogenesis, cell division and differentiation are halted, and the seed enters the maturation phase of its development. Large scale physiological and metabolic shifts accompany the onset of maturation. While the early stages of maturation are characterized by the accumulation of nutrient reserves such as starch, oils, and storage proteins, during late maturation, seeds begin to gradually lose water and acquire DT and longevity (Dekkers *et al*., 2015; Leprince *et al*., 2016; Artur *et al*., 2024). During this maturation drying phase, seeds accumulate protective proteins, non-reducing sugars (e.g. sucrose and raffinose), and ROS-scavenging enzymes (Leprince *et al*., 2016; Kijak & Ratajczak, 2020; Smolikova *et al*., 2020). By the end of maturation drying, orthodox seeds reach their lowest water content, typically between 5-10% relative water content (RWC), and achieve maximum storability. Thus, the temporal regulation of protective mechanisms during late maturation prepares the seed to be stored in the dry state for prolonged periods until conditions are favourable for germination.

The transition of seeds into maturation and the consequent acquisition of DT and longevity are transcriptionally regulated by a complex and yet robust gene regulatory network (GRN) controlled by the hormone abscisic acid (ABA) (Angelovici *et al*., 2010a; Gazzarrini & Song, 2024) and four transcriptional master regulator genes - *LEAFY COTYLEDON 1* and *2* (*LEC1* and *LEC2*), *ABSCISIC ACID INSENSITIVE 3* (*ABI3*), *FUSCA 3* (*FUS3*), often termed the LAFL network (Angelovici *et al*., 2010a; Gazzarrini & Song, 2024). The LAFL network is directly involved in the transcriptional regulation of genes that encode for protective proteins such as Late Embryogenesis Abundant proteins (LEAs) and Heat Shock Proteins (HSPs) (Verdier *et al*., 2013; González-Morales *et al*., 2016; Leprince *et al*., 2016). Interestingly, several transcripts expressed during maturation are either translated immediately into proteins or are translated at later maturation timepoints, during seed drying, pointing towards a role of translational regulation during seed maturation (Chatelain *et al*., 2012; Verdier *et al*., 2013; Sajeev *et al*., 2019; Bai *et al*., 2020, 2023). While the transcriptional regulation of seed maturation is well established, far less is known about the contribution of post-transcriptional regulation to this process. Over the last decade, substantial progress has demonstrated that plant development and stress responses are controlled not only at the transcriptional level but also through multiple post-transcriptional mechanisms (Prall *et al*., 2019). This is particularly evident during seed germination, a stage at which the regulation of mRNA fate—through control of mRNA stability and translation—plays a critical role in the timing and success of germination (Sajeev *et al*., 2022; Guo *et al*., 2023a; Bai *et al*., 2026). In the recent years, a specific mRNA degradation pathway called co-translational mRNA decay (CTRD) has been established as a wide-spread mechanism through which nuclear mRNAs are degraded while still being translated (Pelechano *et al*., 2015; Carpentier *et al*., 2024). CTRD was demonstrated to allow plant cells to rapidly adjust the active pool of mRNA accessible for translation in response to developmental and environmental cues (Carpentier *et al*., 2020; Guo *et al*., 2023b; Zhang *et al*., 2023; Dannfald *et al*., 2025; Deragon & Merret, 2025). However, little is known about the CTRD status of seeds, especially during seed maturation and drying.

The drying rate of seeds during maturation has a considerable impact on their physiological properties. For example, in soybean (*Glycine max* L.), seed quality traits were better preserved in maturing seeds detached from the mother plant that were slow-dried either within pods or outside the pod under controlled humidity, compared to seeds fast-dried at low humidity (Adams *et al*., 1983; Blackman *et al*., 1992; Sinnecker *et al*., 2005). In common beans (*Phaseolus vulgaris* L.), slow drying seeds outside the plant during early maturation enhanced DT (Sanhewe & Ellis, 1996). Similarly, in foxglove (*Digitalis purpurea* L.), placing maturing seeds in near-saturated environments or pre-drying under high-to moderate-humidity conditions prior to desiccation improved DT (Hay, 1995). Similar observations of quality improvement by slow drying inside or outside pods in orthodox seeds have been reported for several other species (Ooms *et al*., 1993; Corbineau *et al*., 2000; Samarah, 2006; Samarah *et al*., 2010). A notable share of seeds at harvest are premature due to the inability to complete maturation drying on the mother plant. This leads to poor-quality seed lots that lack proper storability and vigor. Varying drying regimes have also been shown to play an important role in improving the quality of seeds after priming treatments (Butler *et al*., 2009; Fabrissin *et al*., 2021; Veser *et al*., 2025). This mounting evidence suggests that the drying rate and the ABA-pathway act as two major directive forces enabling the temporal regulation of seed maturation, which leads to enhanced seed quality traits and DT. However, a comprehensive understanding of the molecular mechanisms underlying this drying-induced physiological effect in seeds is still lacking.

In this study, we investigated the effects of both fast and slow drying regimes on maturing seeds in the model plant *Arabidopsis thaliana* L. We examined the physiological impact of drying seeds either inside or outside the fruits (siliques). Additionally, we analyzed the hormonal, transcriptional, post-transcriptional, and translational changes in seeds resulting from these different drying regimes. To further explore the role of ABA levels in the physiological and transcriptional changes induced by drying, we utilized the ABA deficient mutant *aba2-1*. Our findings show that maturing seeds can sense and respond to different drying regimes characterized by distinct transcriptomic signatures. Slowing drying had a more profound impact on both seed physiology, transcriptional, post-transcriptional and translational landscape of seeds compared fast drying. These findings have the potential to help develop more effective post-harvest and post-priming drying strategies. Furthermore, this research may lead to the identification of molecular markers to monitor the drying process. Overall, this study enhances our understanding of the relationship between drying strategies and the molecular dynamics in maturing seeds.

## Results

### Slow drying accelerates desiccation tolerance induction during seed maturation

To study the molecular effects of different drying regimes on seed maturation in *Arabidopsis*, we first determined the timing of DT acquisition, a physiological trait tightly linked with water loss and the foundation of seed longevity. The development of *Arabidopsis* seeds spans approximately 20-26 days, depending on the growth conditions, with the acquisition of DT starting at 12 days after pollination (DAP) and fully established by 16-18 DAP (Baud *et al*., 2002; González-Morales *et al*., 2016; Jing *et al*., 2018; Artur *et al*., 2024). We reassessed DT acquisition using a previously established method where seeds are removed from the fruit (siliques) and subjected to fast drying (FD) in a drying cabinet (30% relative humidity (RH) at 22°C in the dark) for 2 days (Ooms *et al*., 1993, 1994). Similar to previous reports, we found that DT is acquired between 12-16 DAP (Fig. 1a), with almost all seeds having DT by 16 DAP. In parallel, the water content of the seeds decreased sharply between 10-12 DAP but maintained a steady decline until plateauing at 22 DAP (Figs. 1a and Fig. S1). The seed dry weight, on the other hand, increased until 16 DAP and then plateaued, indicating the end of seed filling and the onset of late maturation drying (Fig. S1).

**Figure 1.**
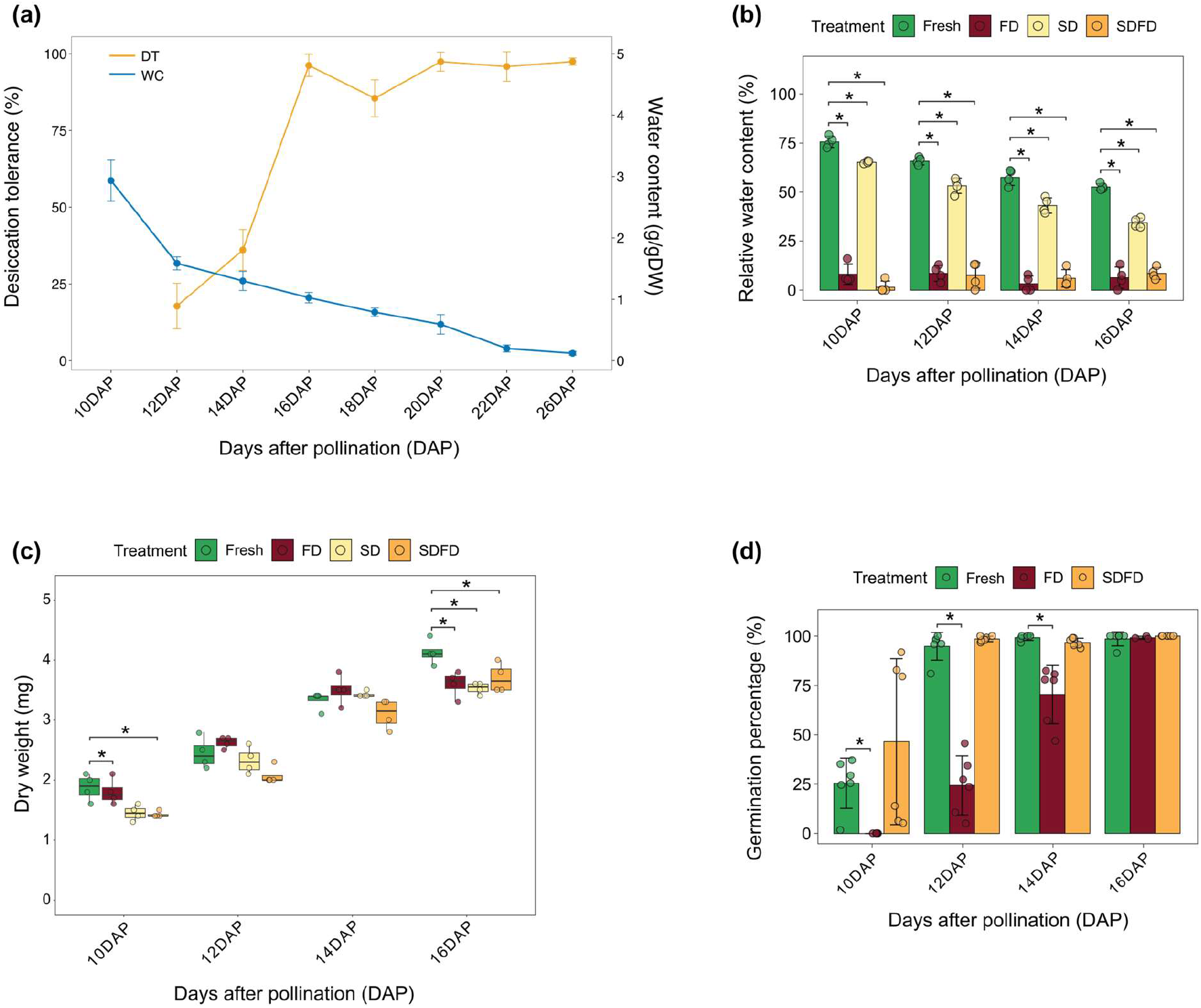
Physiology of maturing *Arabidopsis thaliana* (Col-0) seeds. (**a**) Acquisition of desiccation tolerance and water content change in maturing seeds. Desiccation tolerance is expressed as the percentage of seeds that survive (i.e., germinate) following fast drying at 30% RH and 22°C for 2 days. Water content is expressed as grams of water per gram of dry weight (g/g DW). Means and standard deviation of four replicates (n=4) are presented. (**b**) Changes in relative water content (RWC) and (**c**) changes in dry weight of maturing seeds of 10, 12, 14, and 16 DAP subjected to no drying (Fresh), 2 days fast drying (FD), 2 days slow drying (SD), and 2 days SD followed by 2 days FD (SDFD). Error bars in (b) indicate standard deviation. In both (b) and (c), four replicates (n=4) were used for each time point and treatment combination. (**d**) Mean germination capacity (%) of maturing seeds subjected to different drying regimes - no drying (Fresh), FD, and SDFD. Error bars indicate standard deviation calculated from six replicates (n=6). Statistical significance of treatment effect in (**b**) and (**d**) was calculated using the Kruskal-Wallis test followed by pairwise comparison using the Wilcoxon rank-sum test. For (**c**), significance was determined using one-way ANOVA followed by pairwise comparison using a t-test. Asterix (*) indicates a significant difference of FD, SD, and SDFD samples from Fresh samples of the same time point (*p-value < 0*.*05*).

Due to the dynamic changes taking place during early time points of seed maturation with respect to DT and water loss, we selected 10, 12, 14, and 16 DAP to assess the effect of different drying regimes on seed maturation by primarily focusing on DT acquisition. We altered the natural drying process of these premature seeds by removing them from the mother plant and subjecting them to different drying regimes, where each regime allowed the seeds to dry at different speeds. We applied three drying treatments – i. the standard FD method used to phenotype for DT acquisition (fast drying regime), ii. slow drying (SD) seeds inside the fruit (silique) tissue in a drying cabinet (30% RH at 22°C in dark) for 2 days (slow drying regime), and iii. 2 days of SD followed by 2 days of FD (slow-fast drying regime).

We opted for silique drying as the method for imposing SD since for small seeds (like in *Arabidopsis*), the fruit can act as a barrier reducing water loss as demonstrated in previous reports (Adams *et al*., 1983; Ooms *et al*., 1993; Corbineau *et al*., 2000; Samarah *et al*., 2010).

We investigated the effects of these three drying regimes on maturing seeds at different physiological levels. Both FD and SD caused a significant change in the relative water content (RWC) of the seeds, albeit at different magnitudes. FD reduced RWC to below 10% at all time points (Fig. 1b). On the contrary, 2 days of SD led to less drastic (yet significant) changes in water levels and resembled the natural RWC change observed over 2 days during *in planta* maturation (Fig. 1b). For example, the RWC change observed between 12 DAP Fresh and 12 DAP SD seeds was comparable to the RWC change between 12 DAP Fresh and 14 DAP Fresh seeds. As expected, the SDFD treatment led to a similar decrease in RWC as FD. However, unlike RWC, the dry weight of the seeds was largely unaffected by the drying regimes. The only notable changes were seen at 10 and 16 DAP where the treatments led to reduced dry weight (Fig. 1c). However, no instance of dry matter gain was observed, even during SD when seeds were still attached to the fruit tissue. On the other hand, similar to changes in RWC, DT acquisition between 10 – 16 DAP was significantly influenced by the drying regime. We compared seed germination between untreated seeds (Fresh), and desiccated seeds via FD and SDFD (Fig. 1d). SD prior to FD (i.e., SDFD) induced DT in seeds as early as 10 DAP, albeit with high variation (Fig. 1d). At 12 DAP, all SDFD-treated seeds were desiccation-tolerant, in contrast to *in planta* DT maxima that were only obtained at 16 DAP (Fig. 1a, d).

Since ABA plays a major role in the onset of seed maturation, we wanted to understand the ABA dynamics following the three drying regimes and its association with the observed physiological changes. To investigate this, we measured ABA levels in both seed and silique tissues. ABA content in freshly harvested seeds from 10 – 16 DAP showed a bell-shaped pattern with a peak at 12 DAP and lowest amount in mature dry seeds (DS) (Fig S2a). SD accelerated this bell-shaped trajectory of ABA dynamics. For instance, 10 DAP SD seeds already showed an ABA levels similar to 12 DAP Fresh seeds and there was sharp decline in ABA content in 12 DAP SD seeds which resembled ABA levels of 16 DAP Fresh seeds. In contrast, FD either led to no changes in ABA levels (10 and 12 DAP) or led to significantly lower levels (14 and 16 DAP) compared to Fresh seeds. In case of both SD and FD, the seed ABA level showed a time point-dependent rather than a drying treatment-dependent change. However, an opposite trend was visible for the ABA content in silique tissues. In fresh silique tissues the ABA content maintained steady levels between 10 – 16 DAP but showed a high peak in mature dry siliques. Interestingly, ABA levels were consistently significantly higher under SD in all maturation timepoints (Fig. S2b) whereas FD only showed significant increase at 14 DAP. Overall, the disparity in the seed and silique ABA content may be due to limited transfer of ABA between the tissues due to the drying treatments.

### Global gene downregulation accompanies the acceleration of DT induction and ABA dynamics during slow drying

The molecular mechanisms that govern how seeds respond to different drying regimes remain poorly understood, particularly regarding how they perceive water loss. To address this gap, we investigated the transcriptome by performing RNA-seq on seeds of 10 – 16 DAP subjected to no drying (Fresh) or any of the three drying regimes (FD, SD and FDSD) (Fig. 2a). We also included mature dry seeds (26 DAP) in the RNA-seq as a reference for the transcriptome of *in planta* matured dry seeds (DS). All the experiments were performed using three biological replicates. Our goal was to determine whether seeds can detect varying rates of water loss and initiate a corresponding transcriptional response. Furthermore, we aimed to determine whether these transcriptional responses could explain the early acquisition of DT and the accelerated ABA dynamics in seeds subjected to a slow-drying regime.

**Figure 2.**
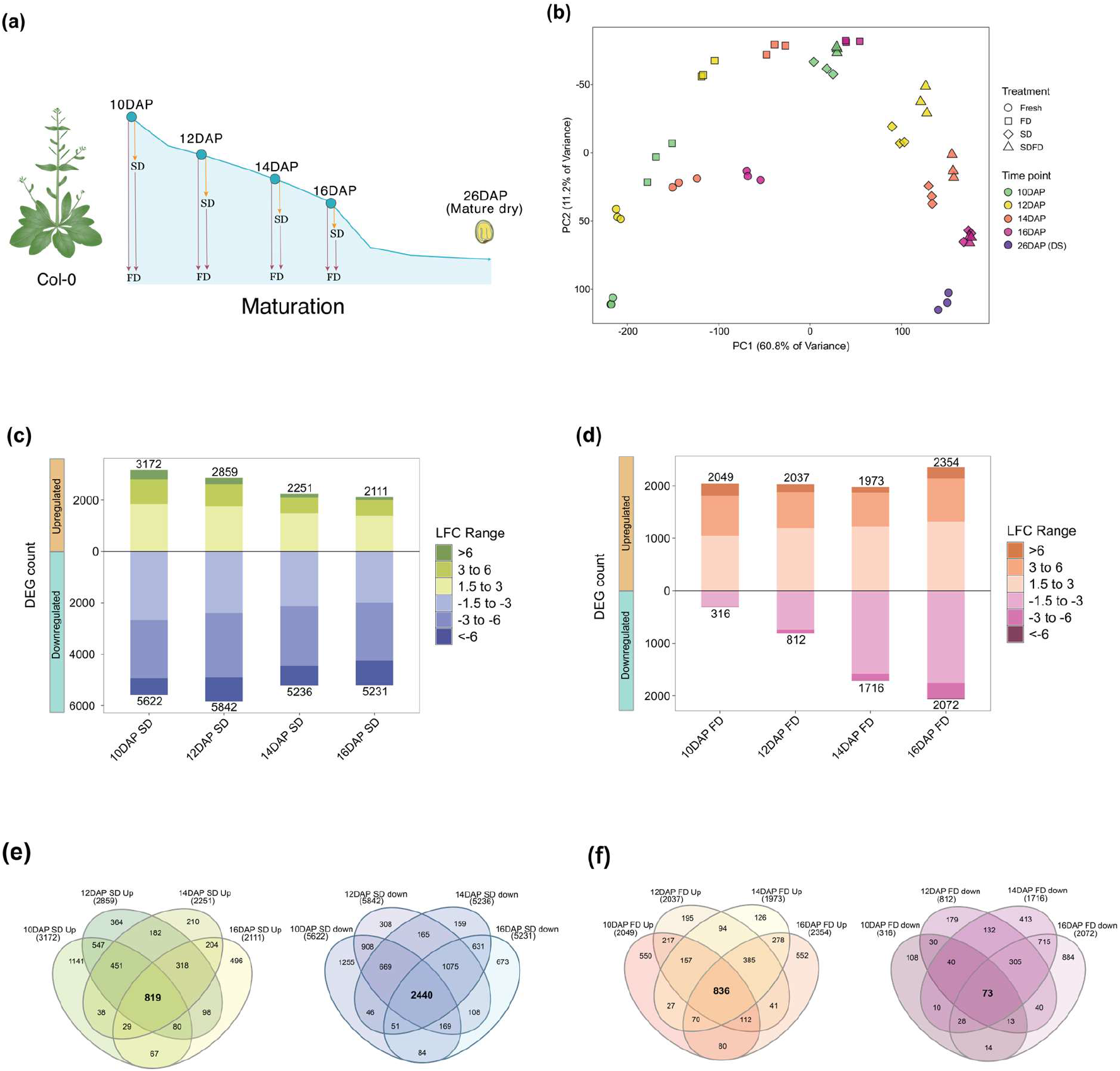
Transcriptional response of maturing Col-0 seeds to different drying rates. (**a**) Design of the bulk RNA-seq experiment. Samples include maturing seeds of 10, 12, 14, and 16 DAP that are either freshly harvested (Fresh) or treated with FD, SD, and SDFD. Mature dry seeds (26 DAP) were included as a reference for complete seed maturation. (**b**) Principal component analysis (PCA) showing the distribution of the RNA-seq samples. Shapes and colors indicate the treatments and time points, respectively. (**c**) Number of differentially expressed genes (DEGs) that are up- or downregulated in SD samples compared to Fresh samples for each time point. Counts include genes that have a significance of *p-adj < 0*.*01* with a Log2 Fold Change (LFC) of > 1.5 or <-1.5. (**d**) Number of differentially expressed genes (DEGs) that are up- or downregulated in FD samples compared to Fresh samples for each time point. Counts include genes that have a significance of *p-adj < 0*.*01* with a LFC of > 1.5 or <-1.5. (**e**) Venn diagram showing core up (left) or downregulated (right) genes at all time points after SD treatment. (**f**) Venn diagram showing core up (left) or downregulated (right) genes at all time points after FD treatment. Numbers between parenthesis indicate the total number of DEGs in each sample (timepoint x treatment).

Initial exploration of the RNA-seq data using principal component analysis (PCA) revealed a good reproducibility between the biological replicates and indicated that PC1 and PC2 together accounted for 72% of the variation observed in the transcriptome, with PC1 explaining the majority at 60.8% (Fig. 2b). According to the PCA, all drying regimes i.e., FD, SD, and SDFD led to a shift in the transcriptional state of the seed at all time points. However, the changes induced by SD and SDFD were more pronounced than those caused by FD. Both SD and SDFD caused large shifts along PC1, towards DS.

Although this shift was consistent for all time points, more mature seeds (such as 16 DAP submitted to SD and SDFD) showed greater proximity to DS. A similar trend was evident following hierarchical clustering of RNA-seq samples based on Euclidean distance, where all SD and SDFD samples formed a large cluster alongside DS (indicated as 26 DAP, Fig. S3, green box). This clearly demonstrates that 2 days of SD and 4 days of SDFD accelerated the seed transcriptome, reaching a state comparable to that of DS. This acceleration in the transcriptome aligns nicely with the observed early acquisition of DT (Fig. 1d) and ABA dynamics following SD (Fig. S2a).

To dive deeper into the genes modulating this transcriptional shift, we performed pairwise comparisons between seeds that underwent FD, SD, or SDFD against the untreated seeds (Fresh) of the same time point to identify differentially expressed genes (DEGs) (Log_2_ Fold Change (LFC) > 1.5 or < -1.5 and p-adj < 0.01). All three drying regimes led to the upregulation of more than 2,000 genes across all time points (Fig. 2c, d and Fig. S4a). In contrast, the trend of downregulated genes varied between the fast-drying regime (i.e., FD) and the two slow-drying regimes (i.e., SD and SDFD) (Fig. 2c, d, and Fig. S4a). For SD and SDFD, a large number of genes (over 5,000) were downregulated at all time points. However, this was not the case for FD samples, where the number of downregulated genes increased progressively with maturation time, reaching a peak at 16 DAP with 2,072 downregulated genes. This progressive increase in downregulated genes under FD coincided with the increase in DT, i.e., germination (%) following desiccation between 10-16 DAP, and was opposite to the moisture content pattern of the seeds during this time (Fig. 1d and Fig. S1).

To identify the core genes that were differentially expressed under each specific drying regime, regardless of maturation time point, we analyzed the overlap of upregulated and downregulated DEGs among all time points (Table S1). A total of 836, 819, and 1,143 core genes were consistently upregulated under FD, SD, and SDFD, respectively, at all maturation time points (Fig. 2e, f and Fig. S4b). On the other hand, 2,440 and 2,649 genes were identified as the core downregulated genes under SD and SDFD, respectively (Fig. 2e and Fig. S4b). Interestingly, only 73 genes were identified as core downregulated genes under FD, owing to substantial variation in DEG counts across time points (Fig. 2d). We observed that, on one hand, there is an increase in gene downregulation during DT induction, while on the other hand, SD and SDFD accelerate both transcriptome changes and DT acquisition (Fig. 1d and Fig. 2b). Based on this, we hypothesized that this global downregulation of genes is accelerated under slower drying regimes. To investigate this, we examined the overlap between the core SD downregulated genes and the FD downregulated genes at each time point. Indeed, we found that the overlap between the FD downregulated genes at each time point and the core SD downregulated genes increased with the progression of maturation (Fig. S5a). This gradual increase in the number of shared downregulated genes between FD and SD core genes followed a similar trend as the increase in DT and was opposite to the moisture level of the seeds during maturation (Fig. S5a and Fig. 1d). On the other hand, the overlap between the core SD upregulated genes and FD upregulated genes at each timepoint was relatively consistent (Fig. S5b). Together, these findings suggest that global gene downregulation accounts for the accelerated induction of DT by SD, which could result not only from changes in moisture content but also from alterations in ABA levels in the seeds.

### Distinct transcriptome signatures underlie seed responses to different drying regimes

The core up- or downregulated genes might reflect consistent transcriptome responses of seeds to specific drying regimes. To identify unique responses to each drying regime (drying regime-specific genes) and differentiate them from common responses across different drying regimes (general drying-related genes), we investigated the overlap between the core up and downregulated genes of each drying regime. The largest overlap was found between the core SD and SDFD downregulated genes, with 1,984 genes being downregulated exclusively during these two drying regimes (Fig. 3a and Table S2). These genes (labelled as “Shared SDFD-SD down”) were enriched in GO terms like photosynthesis, response to light, carbohydrate metabolisms, fatty acid biosynthesis, and several other terms that related to primary and secondary metabolism, cell growth, and seed development (Fig. 3b). In contrast, the exclusive overlap between FD and SDFD downregulated genes (“Shared FD-SDFD down”) was minimal (8 genes) due to the small number of core FD downregulated genes (73 genes, Fig. 2f). An additional 56 genes were commonly downregulated under all three drying regimes (“Shared SDFD-FD-SD down”) (Table S2). Interestingly, 600 genes were downregulated only under SDFD. These genes were enriched in unique categories such as cutin biosynthesis, mucilage biosynthesis, beta-D-glucan biosynthesis, cellular response to ABA, amino acid transport, and megasporogenesis. This shows that the combination of SD followed by FD resulted in the downregulation of specific biological processes that were not observed during either SD or FD alone. On the other hand, the 394 genes specific to SD (“SD down”) showed a unique enrichment for GO terms related to leaf development, auxin transport, cell division and differentiation, cuticle development, xylem-phloem patterning, and hyperosmotic salinity response (Fig. 3b). It is intriguing that these 394 genes are specific to SD and not shared with SDFD since both of them share the first 2 days of SD. This suggests that their expression is likely altered during the subsequent 2 days of FD, indicating that these genes can be modulated by distinct drying regimes.

**Figure 3.**
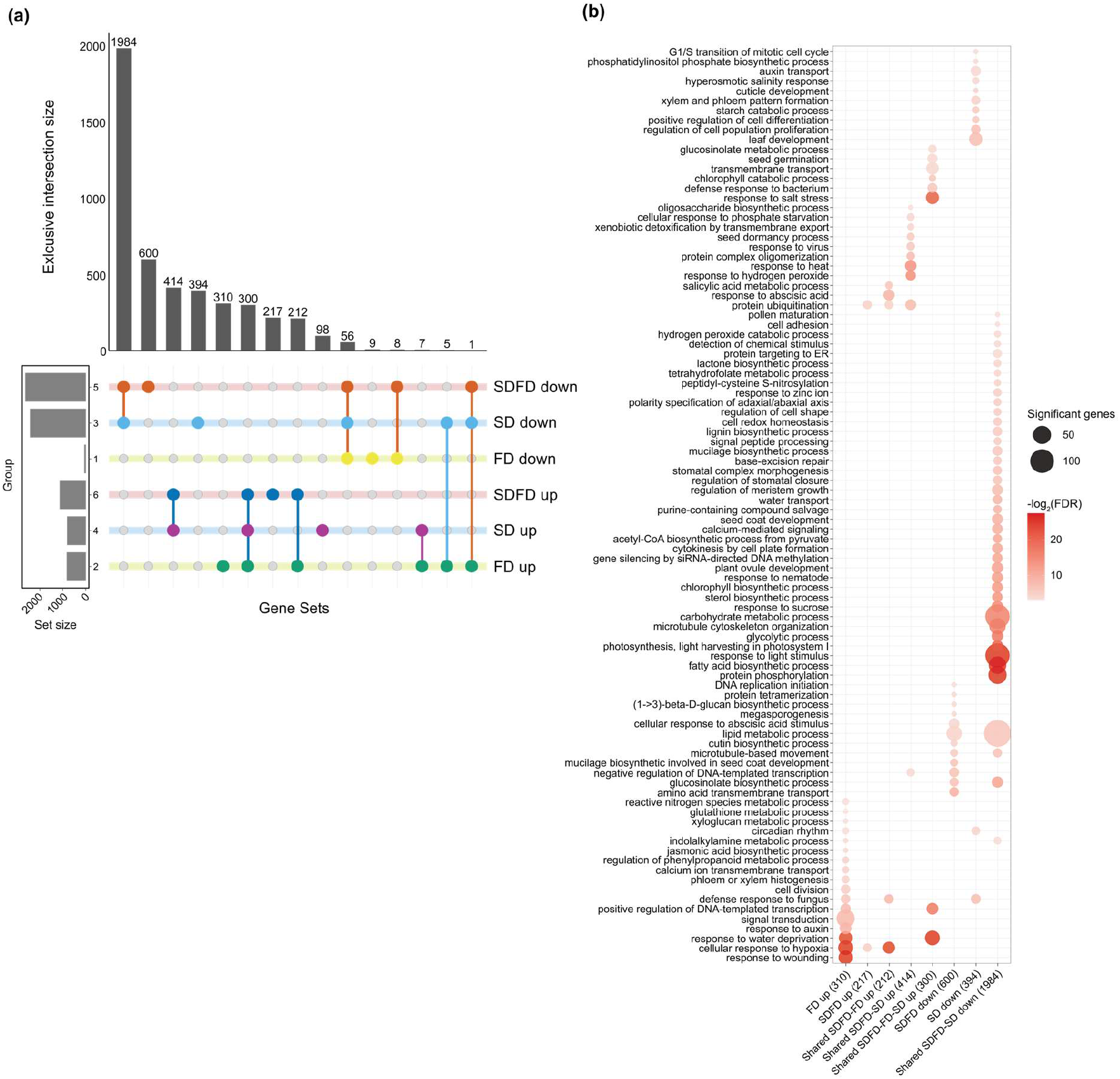
Overlap between core FD, SD, and SDFD DEGs. (**a**) Upset plot showing overlap in core DEGs following FD, SD, and SDFD. (**b**) Gene ontology (GO) enrichment of core DEGs that are unique to each or shared between the drying regimes as shown in (a). Color intensity indicates the -log2(FDR) value of the GO term. The size of the circles corresponds to the number of significant genes belonging to that GO term in the target gene list. Only gene sets that had sufficient number of genes to perform GO enrichment are shown.

The core upregulated genes showed an overlap of 300 genes across FD, SD, and SDFD (“Shared SDFD-FD-SD up”) (Table S2). The lack of specificity in relation to the speed of drying emphasizes that the activation of these genes is independent of the drying regime (Fig. 3a). These shared core upregulated genes were enriched in genes related to response to water deprivation, salt stress, defense response to bacterium, seed germination, chlorophyll catabolism, and glucosinolate metabolism (Fig. 3b). Another 414 genes were shared only between SDFD and SD (“Shared SDFD-SD up”) which were enriched for oligosaccharide biosynthesis, response to heat, hydrogen peroxide, phosphate starvation, seed dormancy, and protein complex oligomerization (Fig. 3b). These genes are activated under a slow-drying regime but unaffected by a subsequent fast drying (in SDFD). On the contrary, the 212 shared genes between SDFD and FD (“Shared SDFD-FD up”) represent genes that are commonly induced under a fast-drying regime. These genes showed an enrichment for response to hypoxia, abscisic acid, defense response to fungi, salicylic acid metabolism, and protein ubiquitination. Notably, an additional set of 310 and 217 genes were unique to FD and SDFD, respectively (Fig. 3a). The 217 “SDFD up” genes were only enriched for response to hypoxia and protein ubiquitination. These genes might require both SD and FD for activation. On the other hand, the 310 “FD up” genes showed a clear enrichment for several biotic and abiotic stress responses such as responses to wounding, fungi, hypoxia, and water deprivation, jasmonic acid (JA) biosynthesis, metabolism of glutathione, indolalkylamine, phenylpropanoid, and xyloglucan (Fig. 3b). These processes are related to activation of a defense response and defense related metabolites (Fonseca *et al*., 2009; Hasanuzzaman *et al*., 2017; Dong & Lin, 2021; Sun *et al*., 2022). This strongly suggests that the FD-specific response is dominated by defense-related transcriptional activity and could indicate potential structural cellular damages induced by the rapid loss of water from premature seeds.

### Co-expression network analysis reveals gene modules specific to fast and slow drying

The distinct and shared transcriptional responses of seeds to various drying regimes led us to investigate the gene regulatory networks (GRNs) associated with the drying-regulated genes. To increase the resolution and coverage of our analysis, and to understand the behaviour of gene modules under both normal maturation (when seed quality traits are acquired) and during different drying regimes, we combined the RNA-seq data from this study (Fresh, FD, SD, and SDFD samples) with the RNA-seq data from the transcriptome atlas of *Arabidopsis* seed maturation (Fresh seeds samples from 12 – 26 DAP at 2 days interval) (Artur *et al*., 2024). This was feasible due to the similarity in plant growth conditions and the high similarity of time points (12, 14, 16, and 26 DAP Fresh) between the two studies, according to PCA (Fig. S6). This combined approach allows to bridge GRN of seed maturation and of seed water loss.

We identified a total of 14 gene modules (M1-M14), each showing dynamic behavior during maturation and drying (Fig. 4a, b, and Table S3). Interestingly, several modules deviated from their general pattern during standard maturation time-course (10 – 26 DAP), specifically under fast (FD) or slow (SD and SDFD) drying (e.g., M3, M6, M8, M9, M10, M14) (Fig. 4a, b). On the other hand, some modules showed no overall change in their patterns between the standard maturation and the drying regimes (e.g. modules M1, M2, M4, M5, M7, M11, M12, M13). These fourteen distinct modules were enriched in a diverse range of GO terms, some of which were shared amongst multiple modules (e.g. response to abscisic acid, water deprivation, salt stress, hypoxia and wounding) (Fig. S7, marked red), while several others were unique to each module (see Table S4 for full list of GOs). To identify drying regime-specific gene modules and their association with the physiological traits, we performed module-trait correlation analysis (Fig. 4c and Table S5). Based on the dynamics of the modules during maturation, under drying-regimes, correlation with physiological traits, and GO enrichment, we decided to focus on the two modules M5 and M6. M5 genes showed a net positive z-score during late maturation and also during FD and SD. However, the positive trend was much higher during SD compared to FD. Interestingly, M5 showed the most significant positive correlation with DT germination (%). Moreover, M5 also showed significant positive correlation with ABA content in siliques, and negative correlation with RWC and with ABA content in seeds (%) (Fig. 4c). Overall, M5 genes were associated with response to water deprivation, oxidative stress, salt stress, protein ubiquitination, seed dormancy, and chloroplast organization (Fig. S7). This indicates that the genes in the M5 module may influence drying responses during seed maturation. Indeed, the GRN from the M5 gene module included major TFs previously reported to regulate DT, such as *ABI5 (AT2G36270), PLATZ2 (AT1G76590)*, and *RAP2*.*1 (AT1G46768)* (Table S6). Interestingly, most genes in this GRN were upregulated under SD, while a large part of it was downregulated under FD (Fig. 4d and Fig. S8a). This differential response of M5 module genes to FD and SD may underlie the different effects of these two drying methods on the acceleration of seed maturation and acquisition of DT. Thus, M5 genes may reflect part of the core transcriptional program of DT acquisition in *Arabidopsis*.

**Figure 4.**
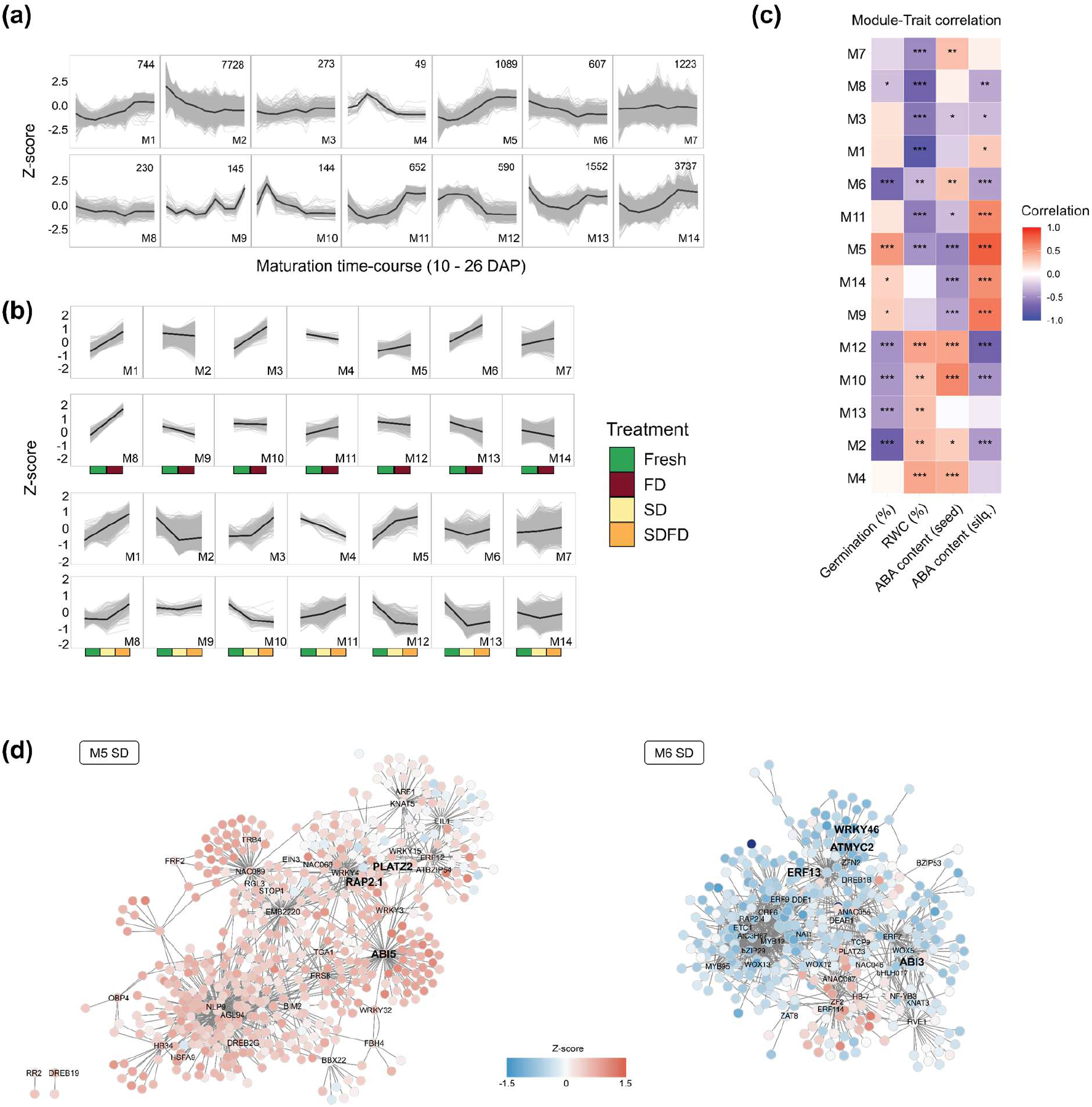
Temporal dynamics of gene modules during seed maturation. (**a**) Changes in z-scores from 14 gene modules during seed maturation. Grey lines indicate z-score of individual genes, and black line indicate the module z-score average. Gene modules were identified based on co-expression using WGCNA by combining the RNA-seq data from this study with the SeedMatExplorer dataset (Artur et al., 2024). (**b**) Dynamics of gene modules under different drying regimes. Colors along the x-axis indicate the treatment. Top two panel rows show the z-score changes of the 14 modules following FD. Bottom two panel rows represent the module z-scores changes after SD and SDFD. For each gene, the z-score was averaged across all time points for the treatment and plotted as grey lines. The black line indicates the module’s mean z-score. (**c**) Module-trait correlation, illustrating the association between each gene module and various physiological traits. Color intensity indicates the strength of the correlation. Asterisks indicate the level of significance (*** - p < 0.0001, ** - p < 0.001, and * - p < 0.01). (**d**) Gene regulatory networks predicted from genes of module M5 (left) and M6 (right) using GRNBoost2. The top 1000 edges were visualized in Cytoscape. The color code indicates the changes in the z-score of gene expression under SD treatment. Nodes corresponding to major TF were shown in each network.

On the other hand, the M6 gene module exhibited a high positive trend under FD but a decreasing trend during standard maturation time-course and SD (Fig. 4b). Moreover, this module showed a strong negative correlation with DT germination (%) (Fig. 4c). M6 was enriched in genes involved in response to wounding, water deprivation, cold, abscisic acid, hypoxia, and oxidative stress (Fig. S7 and Table S4). GRN constructed using the top 1000 connections from the M6 module showed that most genes in the network had a negative z-score following SD (Fig. 4d, right) but a positive Z-score after FD (Fig. S8b). Together, these observations strongly suggest M6 as a candidate gene module containing genes involved in modulating developmental and stress responses to a fast drying regime and may reflect (part of) a stress- or damage response induced during FD. This explains the downward trend of this stress-related module under normal maturation (Fig. 4a).

To test that FD leads to activation of transcriptional programs detrimental for normal maturation; we focused on a set of eight genes. Six of these were selected based on DEG comparisons and consistently showed contrasting expression between FD and SD (always upregulated under FD but downregulated under SD) (Fig. 3a, Fig. S9 and Table S7). The two other genes represented major TF hubs within the M6 module - *ERF13* (*AT2G44840*) and *MYC2* (*AT1G32640*) (Table S6) (Dombrecht *et al*., 2007; Wang *et al*., 2019; Zander *et al*., 2020). Moreover, MYC2 is a known TF in the JA-pathway, allowing us to validate the role of JA-pathways in seed responses to drying. Initially, these mutants showed no apparent abnormality in DT acquisition between 12 and 16 DAP (data not shown), and all produced dry seeds at the end of maturation. Given that DT acquisition is a robust process and tightly regulated, we opted for assessing the longevity of these mutants, which is a more cumulative trait that builds upon DT (González-Morales *et al*., 2016; Leprince *et al*., 2016). Interestingly, four of the eight mutants, namely *aos, erf13, myc2*, and *xth22*, had significantly higher longevity (p50) compared to Col-0, visible at around 28-35 days of controlled deterioration test (CDT) (Fig. S10 and Fig. S11). On the contrary, *nudt7* seeds showed significantly lower longevity than Col-0, with the reduction in gMax (%) visible already from 14 days after CDT (Fig. S11). This clearly demonstrated that part of the FD-specific transcriptome is detrimental for seed maturation and storability, explaining why these genes are suppressed during normal maturation and SD.

### A partially ABA-independent pathway regulates seed responses to drying

Together with the accelerated seed maturation and drying-specific transcriptional responses, we also found that SD led to higher accumulation of ABA in silique tissues and accelerated the ABA dynamics in seed tissues (Fig. S2a, b). This raised a key question – how important are ABA levels in determining the drying-mediated responses in seeds? To address this, we performed phenotypic and transcriptome analysis of the *aba2-1* mutant, a genotype with significantly reduced ABA levels compared to wild type (Col-0) (Leon-Kloosterziel *et al*., 1996; Artur *et al*., 2024). Although the *aba2-1* mutant was more sensitive to both drying regimes (FD and SDFD) at 14 DAP when compared to Col-0, the overall pattern in germination (%) was similar between both genotypes i.e., SDFD led to enhanced DT acquisition than FD alone (Fig. 5a). This implied that a large part of the seed response to specific drying regimes was independent of wild-type ABA levels.

**Figure 5.**
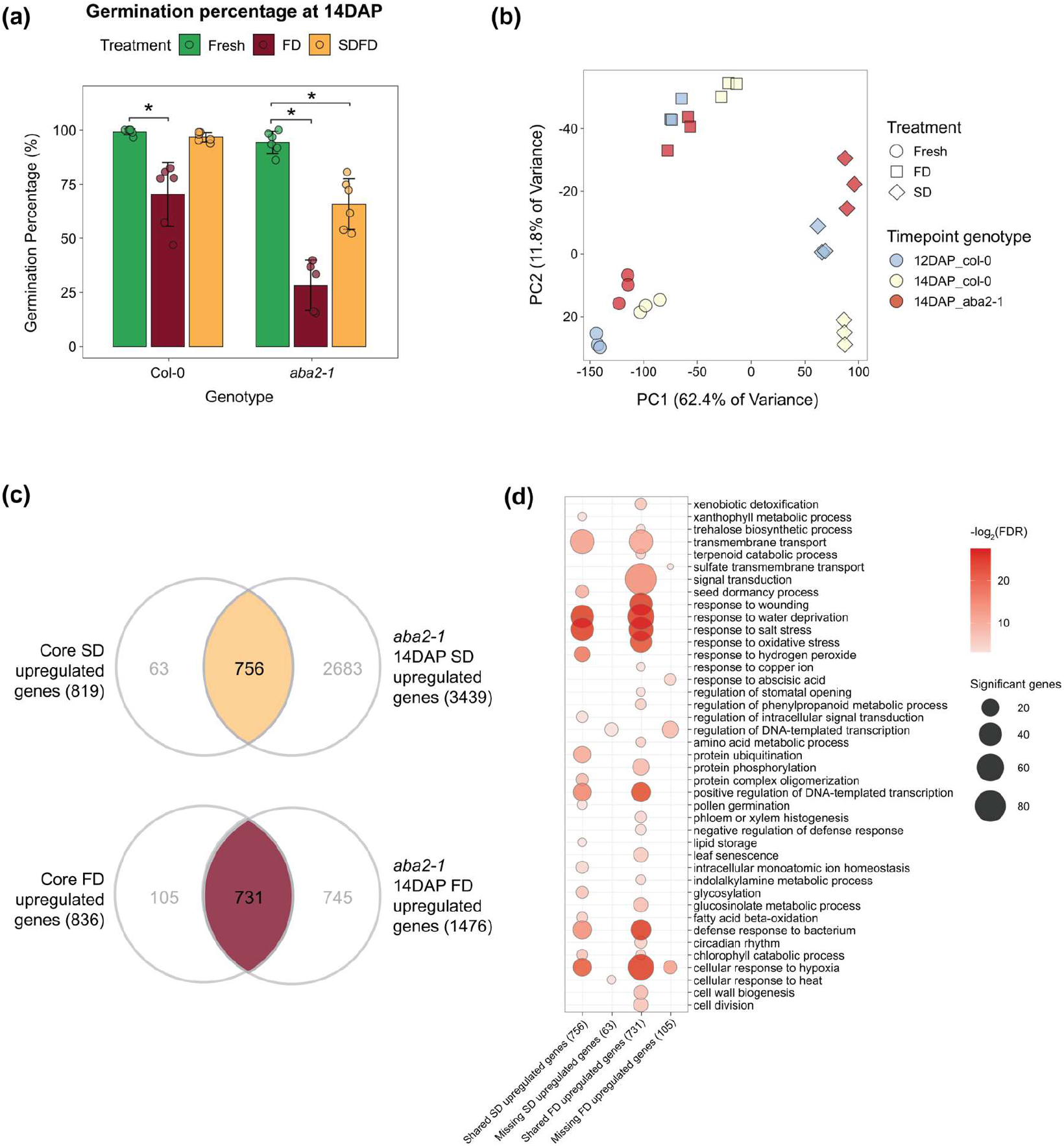
Physiological and transcriptional response of *aba2-1* mutant to different drying rates. Mean germination capacity (%) of 14 DAP Col-0 and *aba2-1* maturing seeds submitted to different drying treatments. Significance was calculated using the Wilcoxon-rank sum test by comparing the FD and SDFD samples to Fresh samples of each genotype. The error bars indicate standard deviation calculated from six replicates (n=6). PCA showing the distribution of 14 DAP *aba2-1* RNA-seq samples with respect to Col-0 samples from 12 DAP and 14 DAP. (**c**) Venn diagram showing the intersection of core FD and SD genes from Figure 2b with those of *aba2-1* 14 DAP. The highlighted part of the Venn diagram indicates core FD and SD genes not upregulated in the *aba2-1* mutant at 14 DAP under the same treatments. (**d**) GO terms enriched in the highlighted genes from (c).

To investigate this, we conducted RNA-seq on seeds from the *aba2-1* mutant at 14 DAP that were freshly harvested and exposed to either FD or SD. We compared the 14 DAP RNA-seq samples from the *aba2-1* mutant to both 12 DAP and 14 DAP Col-0 samples, assessing untreated (Fresh), FD, and SD conditions using PCA (Fig. 5b). Overall, we observed that the *aba2-1* samples followed a similar trajectory as the Col-0 samples, where SD led to a stronger transcriptional shift than FD. 14 DAP Fresh *aba2-1* seeds clustered between 12 and 14 DAP Col-0 Fresh seeds, indicating a developmental delay, which aligned with previous reports (Cheng *et al*., 2014). However, the 14 DAP FD-treated *aba2-1* samples clustered closer to that of 12 DAP Col-0 FD seeds (Fig. 5b), which explained the more sensitive behaviour of these seeds following FD similar to the germination behaviour of Col-0 12 DAP FD-treated seeds (Fig. 1d). On the other hand, 14 DAP SD-treated *aba2-1* samples showed a unique transcriptional state, i.e., they were closer to Col-0 14 DAP-SD samples along PC1 (62.4%) but closer to Col-0 12 DAP-SD seeds along PC2 (11.8%) (Fig. 5b). These distinct transcriptional shifts of the *aba2-1* mutant in response to SD might explain the differences in germination (%) between the mutant and the wild type at 14 DAP (Fig. 1d and Fig. 5a). However, because the overall transcriptomic shift in the mutant followed a similar trend as the wild type (i.e., SD led to greater transcriptomic shift compared to FD), this suggests that much of the drying-mediated phenotypic and transcriptomic responses are largely unaffected under drastically reduced ABA levels.

To identify what genes and biological processes might underly this (partially) ABA-independent response to drying, we compared the core FD and SD DEGs from Col-0 (Fig. 2e, f) with the corresponding DEGs in *aba2-1* (Fig. 5c and Table S8). The reason for using the core genes is that they represent the time point-independent response to drying in the wild type. We used both core up- and downregulated genes for SD, but only the core upregulated genes for FD, since the number of core FD downregulated genes was small (low DEG counts at 10 DAP and 12 DAP). We hypothesized that the shared genes are important for DT, but their response to drying is mainly independent of ABA levels. We found that the majority of the core SD up- and downregulated genes from Col-0 were similarly up- or downregulated in the *aba2-1* mutant (Fig. 5c). More precisely, 756 (out of 819) core SD-upregulated and 2347 (out of 2440) core SD-downregulated genes were present in the *aba2-1* transcriptional response to SD (Fig. 5c and Fig. S12). A similar trend was seen for the 836 core FD-upregulated genes from Col-0 of which 731 were upregulated in the mutant (Fig. 5c). The GO terms response to hypoxia, water deprivation, salt stress, defense response to bacterium, chlorophyll catabolic process, and transmembrane transport were enriched in shared upregulated genes under both FD and SD (Fig. 5d). However, certain biological processes were unique to either FD or SD shared upregulated genes. For instance, responses to wounding, leaf senescence, and oxidative stress were unique to shared FD upregulated genes, while processes related to seed dormancy, protein ubiquitination, pollen germination, lipid storage, and response to hydrogen peroxide were specific to shared SD upregulated genes. We observed a similar trend for shared core SD downregulated genes, where most of the GO terms coincided with those of Col-0 core SDFD-SD down (Fig. S12 and Fig. 3b).

An important aspect of the role of ABA in seed maturation and DT is mediated through the downstream signalling pathway. Two major transcription factors (TFs) involved in this pathway are ABI3 and ABI5. To understand the impact of reduced ABA levels in the *aba2-1* mutant on the downstream canonical ABA signalling pathway, we looked at the expression of *ABI3* and *ABI5* in the Col-0 and in *aba2-1* RNA-seq samples. Notably, these two transcription factors belonged to different gene modules of our GRN. *ABI3* was a member of M6 and exhibited upregulation specifically following FD in Col-0 (Fig. 4d and Fig. S13a). On the other hand, *ABI3* mRNA level was either reduced or unchanged after SD (Fig. S13a). *ABI3* showed a similar expression pattern at 14 DAP in the *aba2-1* background under FD and SD (Fig. S13b). On the other hand, *ABI5* belonged to M5, and its expression increased upon both FD and SD, with an almost two-fold higher induction under SD in Col-0 (Fig. S13a). Interestingly, expression of *ABI5* further increased under SDFD. Basically, the expression of *ABI5* increased linearly as drying time went up from FD to SD to SDFD. This drying-mediated activation of *ABI5* was unchanged in the *aba2-1* mutant (Fig. S13b).

Together, these physiological and transcriptomic analyses indicate that the responses of 14 DAP *aba2-1* seeds to FD and SD closely resemble those of Col-0, with most genes associated with drying-specific responses largely unaffected in *aba2-1*. This indicates that the level of ABA in seeds is not the only factor influencing the activation or repression of downstream genes, such as the transcription factors *ABI3* and *ABI5*. This suggests the existence of candidate signalling pathways that regulate seed responses to drying which operate largely independently of, and potentially parallel to, the ABA-dependent pathway.

### Reduction in co-translational mRNA decay accompanies SD-mediated responses

In addition to transcriptional regulation, we aimed to further explore the impact of SD on post-transcriptional processes, specifically the interaction between mRNA stability and translation during seed maturation. To do so, we focused our analysis on the co-translational mRNA decay pathway (CTRD), which has been shown to participate in many plant processes (Deragon & Merret, 2025). We conducted a 5’P-seq analysis at 12, 14, and 16 DAP under both Fresh and SD conditions. This method enables the simultaneous examination of mRNA decay and ribosome dynamics (Pelechano *et al*., 2015). Indeed, 5′P-seq primarily captures transcripts undergoing co-translational decay. As the 5′-3′ exoribonuclease XRN4 closely follows the last translating ribosome, 5′P reads can be used to infer ribosome footprints and translation activity (Deragon & Merret, 2025). To investigate ribosome dynamics, we first performed a Fast Fourier Transform (FFT) analysis on the different samples (Fig. S14). This analysis revealed a clear three-nucleotide periodicity across all samples, indicative of active ribosome movement and ongoing co-translational decay (CTRD) activity. Next, we performed a meta-transcriptome analysis of 5’P read accumulation around stop codons (Fig. 6a). A clear overaccumulation of 5’P reads at position -16/17 nucleotides (nt) before the stop codons was observed in all samples, a characteristic feature of active CTRD This peak corresponds to the last translating ribosome in a termination step and can be used as a hallmark of CTRD activity (Deragon & Merret, 2025). Interestingly, this peak drastically reduced in seeds of all developmental stages submitted to SD, an indication of reduced CTRD activity (Fig. 6a). As this pathway is fully linked to translation activity, this decrease can reveal a faster shut-down of translation upon SD treatment. Based on previous reports, impaired CTRD at stop positions can lead to an overaccumulation of reads at the start codon, particularly at −13 nucleotides upstream of the start site. This position corresponds to a ribosome stalled in the initiation step, with the ATG codon occupying the P site. To verify this, we checked the meta-transcriptome profile around the start codon of Fresh and SD samples (Fig. 6b). We observed that the metagene profiles of the SD samples around the start codon showed a higher peak at the -13nt position compared to Fresh samples of the same timepoint. To pinpoint the genes causing this pattern, we calculated the read ratio at -13nt between SD and Fresh samples and filtered for genes with a ratio greater than 1 (i.e., higher reads in SD) in at least one timepoint. This resulted in a total of 862 genes that exhibited a higher read accumulation at -13nt in SD samples (Table S9). Removal of these genes from the metagene profile abolished the -13nt peak in both SD and Fresh samples for all time points (Fig. S15a). GO enrichment showed that these genes were involved in processes like response to hypoxia, oxidative stress, water deprivation, abscisic acid, seed oil body biogenesis, lipid storage, seed dormancy, mRNA splicing, and other processes crucial for seed maturation (Fig. S15b). Moreover, when we compared the -13nt peak of these genes to their -16/-17nt peak near the STOP codon within SD samples, a large subset of these genes (n=565) exhibited a higher read accumulation at -13nt START compared to -16/-17nt STOP (ratio > 1) (Table S10). This meant that for these genes, SD led to higher ribosome activity near the START, accompanied by lower activity around the STOP, possibly indicating a faster shutdown of translation.

**Figure 6.**
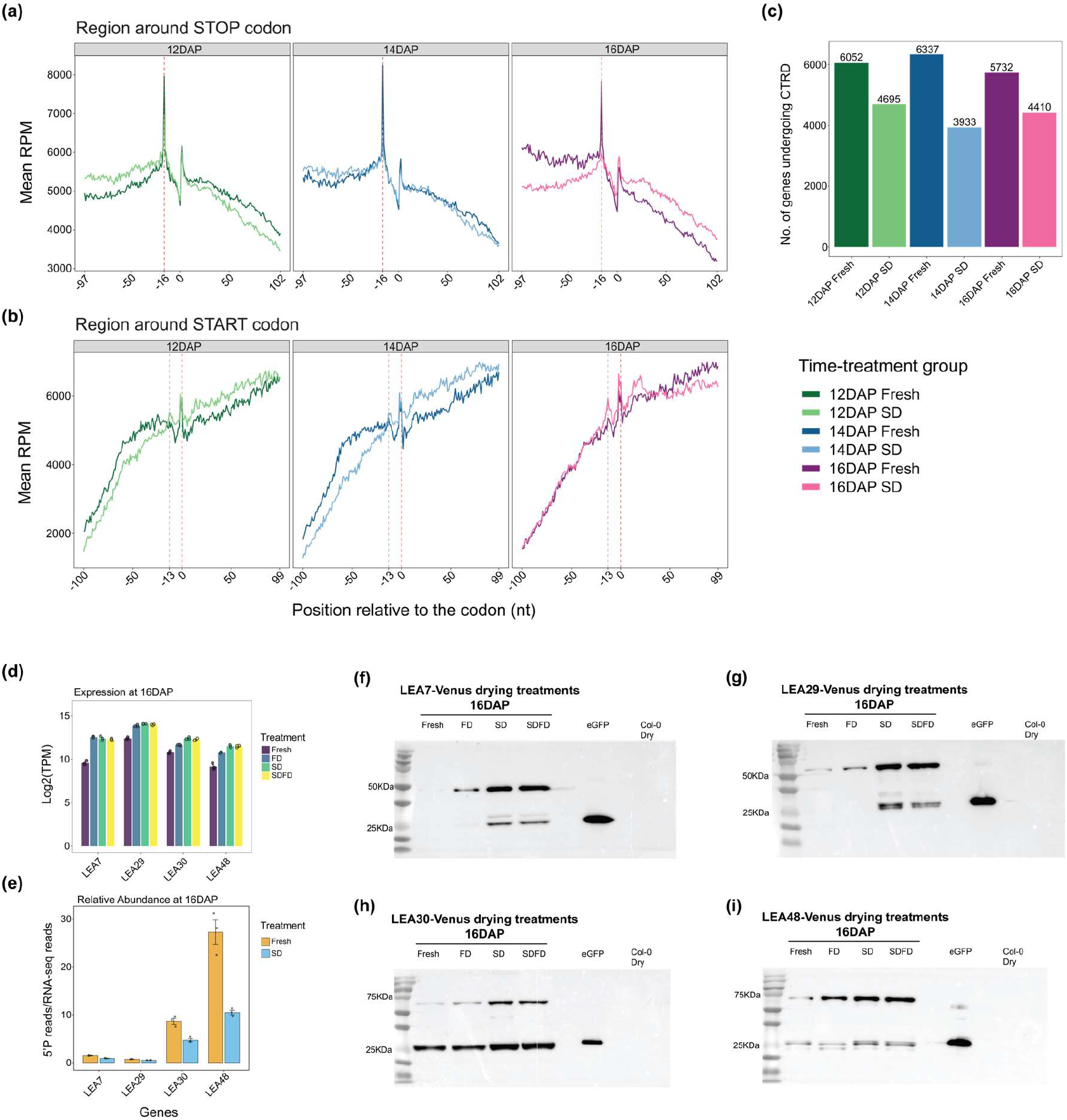
Co-translational decay (CTRD) during seed maturation. (**a**) Metagene analysis showing the accumulation of 5’P reads near the stop codon in Fresh and SD samples of 12, 14, and 16 DAP seeds. Red dashes indicate the stop codon. (**b**) Metagene analysis showing the accumulation of 5’P reads near the start codon in Fresh and SD samples of 12, 14, and 16 DAP seeds. Red dashes indicate the start codon. (**c**) Number of genes undergoing CTRD with a termination stalling index (TSI) value of >= 2. TSI values were calculated by taking the sum of reads at position -16nt and -17 nt and dividing by the mean reads of the flanking 100 nucleotides. (**d**) Transcriptional profile of four *LEA_4* genes – *LEA7* (AT1G52690), *LEA29* (AT3G15670), *LEA30* (AT3G17520), and *LEA4*8 (AT5G44310) based on RNA-seq data at 16 DAP under Fresh, FD, SD, and SDFD treatments. (**e**) Proportion of uncapped transcripts relative to the total RNA pool of the same *LEA_*4 genes at 16 DAP Fresh and SD. Error bars indicate standard error. (**f-i**) Western blot showing the accumulation of each LEA protein in 16 DAP maturing seeds under different drying treatments. Proteins were extracted from *pro::LEA_4:LEA_4-Venus* lines for each *LEA* gene. eGFP and Col-0 mature dry seeds were used as positive and negative controls, respectively.

To further explore CTRD activity in seeds submitted to SD, we calculated the termination stalling index (TSI), which is an indication of CTRD activity on an individual transcript level. Based on a TSI > 2, we found that a high number of genes underwent CTRD during seed maturation (12, 14 and 16 DAP Fresh seeds) (Fig. 6c). Notably, under SD the number of genes undergoing CTRD was lower compared to Fresh samples for all time points. A total of 2,553 transcripts (inclusive intersection) underwent CTRD in all Fresh samples, while 1,546 transcripts underwent CTRD in all SD samples (Fig. S16a). From these, 963 genes were shared between all samples (Fresh and SD) (Fig. S16b). On the other hand, 1,590 and 583 genes underwent CTRD specifically under Fresh and SD treatments, respectively. When we plotted the TSI values of these unique genes between Fresh and SD treatments across different time points, we saw that the TSI values of Fresh-specific CTRD genes were significantly reduced under SD treatments (Fig. S16c). The opposite trend was visible for SD-specific CTRD genes, where the TSI increased under SD (Fig. S16d). CTRD genes unique to Fresh samples were enriched in diverse biological processes such as metabolism (e.g., glycogen metabolism, sucrose transport, mRNA catabolism, etc.), development (e.g., plant ovule development and leaf vascular tissue patterning), and stress (e.g., response to heat, photooxidative stress, hyperosmotic salinity response, abscisic acid, etc.) (Fig. S16f). Conversely, CTRD genes specific to SD were predominantly associated with functions related to stress responses, including response to salt stress, hydrogen peroxide exposure, bacterial defense, and sulfate starvation. Notably, both gene groups showed enrichment in processes related to seed dormancy and chromatin remodeling. This indicated a clear shift in the mRNA pool that was actively undergoing CTRD from Fresh to SD samples. CTRD genes shared between the two conditions showed an intermediate behavior, i.e. the average TSI value of this group of genes was significantly reduced under SD (Fig. S16e) but less pronounced than CTRD genes unique to Fresh samples (Fig. S16c). A similar trend was observed in the metagene profiles of these three gene groups. The metagene profile around the start codon was mostly unchanged between Fresh and SD for all three gene sets. However, there were clear differences in the 16/17nt peak near the stop codons. CTRD genes unique to Fresh samples showed a sharply reduced 16/17nt peak after slow drying (Fig. S17a). In contrast, SD-specific CTRD genes showed almost no change in the -16/17nt peak after slow drying (Fig. S17b). CTRD genes shared between Fresh and SD samples showed a somewhat reduced 16/17nt peak (Fig. S17c). Taken together, transcripts with a high TSI in Fresh samples exhibit reduced CTRD after SD, while the opposite is true for genes with a high TSI following SD. The SD-specific transcripts exhibit increased stalling at the 16/17nt position under SD. As for the 963 genes with a high TSI in both Fresh and SD samples, their CTRD activity is largely unaffected by drying. Overall, these analyses revealed that SD treatment favored translational shutdown during the maturation process.

To finally understand the role of drying on seed maturation and DT, and further explore the relationships between mRNA transcription, translation, and stability with protein accumulation in developing seeds in response to drying, we studied the effect of drying on LEA protein accumulation. In fact, there is a considerable delay between transcripts and proteins LEA accumulation (Bies *et al*., 1998; Verdier *et al*., 2013). However, the post-transcriptional regulation associated with this delay was never assessed. For this purpose, we selected four LEA_4 family genes - *AT1G52690* (*LEA7*), *AT3G15670* (*LEA29*), *AT3G17520* (*LEA30*), and *AT5G44310* (*LEA48*), since these proteins have been shown to be present in seeds (Ginsawaeng *et al*., 2021). The expression of all four genes was upregulated upon both fast (FD) and slow drying (SD and SDFD) treatments, although *LEA30* and *LEA48* had a higher expression under slow drying (Fig. 6d). Interestingly, all four LEAs had a higher relative abundance of 5’P-seq reads in Fresh samples compared to SD samples, again indicating a reduced CTRD activity and suggesting transcripts stabilization under SD (Fig. 6e). To test if this potential mRNA stabilization can have an impact on protein accumulation, we performed western blotting of 16 DAP Fresh, FD, SD, and SDFD-treated seeds of the four LEA_4 proteins using Venus-tagged lines under their native promoter. We selected 16 DAP seeds since by this stage, seeds have acquired DT but still undergo global transcriptional changes under SD and SDFD. Overall, the detection of the different LEA_4 protein accumulation was minimal in 16 DAP freshly harvested seeds (Fig. 6f-i). Although FD led to increased accumulation of the four proteins, especially LEA48 (Fig. 6i), SD and SDFD led to the highest accumulation of all four proteins. Interestingly, the FD and SD treatments had similar effects on *LEA7* and *LEA29* transcription (Fig. 6d), but the effect of SD was more pronounced on protein accumulation levels. This suggests that while FD and SD may enable similar transcriptional activity for these two LEAs, SD might promote greater mRNA stability resulting in higher translational capacity. This elevated protein levels and reduced relative abundance of 5’P-seq reads under SD may indicate that *LEA* transcript stabilization can be an important post-transcriptional process for seed maturation (Fig. 6e).

In summary, our results showed that SD not only reduces mRNA decay activity but also accelerates the shut-down of translation. Interestingly, this was accompanied by a stabilization of LEA transcripts, suggesting a coordinated mechanism. The genes regulated by CTRD are involved in diverse maturation-related processes, including multiple stress-responsive genes, underscoring the essential role of this pathway in seed development and stress adaptation.

## Discussion

Drying is a crucial part of seed development and is necessary for the completion of maturation in orthodox seeds (Kermode, 1990; Angelovici *et al*., 2010b). By drying immature seeds of Arabidopsis under three different drying regimes, we demonstrated that seeds can sense and respond to varying speeds of water loss during maturation. These responses span across multiple layers of regulation, including physiological, hormonal, transcriptional, and post-transcriptional processes. Drying regimes that involved two days of slow drying within the silique had the greatest positive effect on maturation and DT acquisition, consistent with previous findings (Adams *et al*., 1983; Blackman *et al*., 1992; Sanhewe & Ellis, 1996; Sinnecker *et al*., 2005). Strikingly, SD acted as a dual regulator, not only further accelerating maturation but also simultaneously accelerating DT acquisition. This was evident in global transcriptomic patterns, as two days of SD induced a substantially greater shift than two days of maturation *in planta*, possibly by altering the water content status of the seeds but not significantly affecting the dry weight (Fig. 1b, c). This treatment physiologically resembled the late maturation phase, where seeds lose water without further significant increase in dry weight (Fig. S1). By prematurely drying immature seeds in a slow manner, we may have induced a state similar to late maturation. In fact, the transcriptome of SD-treated samples showed a shift towards a mature dry seed (Fig. 1b), further showing that SD accelerated DT acquisition by accelerating the maturation transcriptional program. Another aspect of SD that may have contributed to further maturation is the timeframe allowed for the seeds to continue their maturation-related metabolic processes. In contrast, FD provided a much narrower timeframe before seeds were fully desiccated. This difference is evident in the number of upregulated and downregulated genes under FD, SD, and SDFD. While the number of upregulated DEGs under all three drying regimes was relatively comparable, the number of downregulated DEGs was drastically lower under FD (Fig. 2c, d, and S4a). This highlighted an important aspect about seed responses to drying, which is the difference in timescales between upregulation of desiccation-related processes versus downregulation of related processes. The latter appears to require more time, a gradual rate of water loss, or a combination of both. The significantly lower survival rate (as measured by germination percentage) of immature seeds after FD compared to SDFD may be due to the seeds’ inability to downregulate the necessary genes during FD (Fig. 1d and Fig. 2d). As more genes become downregulated under FD conditions, survival rates increase, peaking at 16 DAP (Fig. 1d, Fig. 2d and S5b). This emphasizes the importance of the often-overlooked aspect of desiccation response: the downregulation of metabolic processes that can be harmful to seeds in a dry state. A study on desiccation-tolerant and intolerant green algae revealed that the key difference between these two groups was the significant downregulation of genes linked to various metabolic processes in tolerant species compared to intolerant ones (Peredo & Cardon, 2020). It appears that SD provides enough time for proper downregulation, while FD leads to a desiccated glassy state before metabolic processes can shut down.

The specific genes that were consistently upregulated under each drying regime (core upregulated genes) shed new light on how seeds respond to drying. Cross-comparisons among the different drying regimes revealed that 300 genes were shared across all three conditions. These genes likely represent those that are generally induced by drying. The majority of these genes, specifically 236 out of 300, belong to modules M1, M5, and M11. Each of these modules exhibited an upward trend during the late maturation stages (Fig. 4a) and across all three drying regimes (Fig. 4b). On the other hand, several genes were unique to each drying regime. Among these, 310 genes that were specifically induced under FD. These genes were enriched in processes related to both biotic and abiotic stresses, including the JA biosynthetic process. This reinforces the role of defense pathways linked to JA in seed maturation (Righetti *et al*., 2015).

Previous studies have noted the involvement of defense-related genes in seed maturation and longevity. Two such genes, *WRKY3* (AT2G03340) and *NFXL1* (AT1G10170), are crucial for the acquisition of seed longevity (Righetti *et al*., 2015; Wang *et al*., 2023). *WRKY3* is categorized within module M5, while *NFXL1* is within module M1 (see Table S3). Both of these modules exhibited an upward trend during late maturation and across all drying regimes, further confirming that these defense-related genes are activated during the drying process. Notably, we identified an unprecedented link between defense-related genes and maturation. Our observations revealed that genes activated specifically during fast drying of immature seeds negatively impact seed longevity. Four of the T-DNA mutants we tested, namely *aos, erf13, myc2*, and *xth22*, showed a significantly higher longevity compared to wild type (Fig. S10). Our findings, in conjunction with previous reports, suggest that defense-related genes can play both positive and negative roles in seed longevity. The speed of drying largely determines this role.

We saw that most of the transcriptional response of seeds to drying was unaffected in the *aba2-1* mutant. The differences that were present were most likely due to the developmental differences between Col-0 and *aba2-1* mutants at the same time point (Cheng *et al*., 2014). This high similarity in induction of drying-related genes between Col-0 and *aba2-1* suggests that many of the genes responsible for seed responses to drying and DT acquisition are not solely dependent on wild-type ABA levels. Likely, an alternative ABA-independent but drying-induced pathway may regulate the GRN controlling DT acquisition. The existence of an ABA-independent osmostress-responsive pathway is well-established (Shinozaki & Yamaguchi-Shinozaki, 2000; Roychoudhury *et al*., 2013; Yoshida *et al*., 2014). Known ABA-independent dehydration-responsive genes such as *DEHYDRATION-RESPONSIVE ELEMENT BINDING PROTEIN 2A* (*DREB2A*, AT5G05410) and *responsive to dehydration 29A* (*RD29A*, AT5G52310) were among the core SD upregulated genes in our study (Table S3). Recent studies have shown that Raf-like mitogen-activated protein kinase kinase kinases (Raf-MAPKKKs) function upstream of both ABA-dependent and independent Snf1-related protein kinase2 (SnRK2s) under osmostress (Takahashi *et al*., 2020; Fàbregas *et al*., 2020). These ABA independent pathways often function in parallel or in conjunction with the canonical ABA signaling pathway. In our study, we found that the expression patterns of two major TFs involved in seed maturation and DT (*ABI3* and *ABI5*) was unchanged in the *aba2-1* mutant upon SD and FD when compared to the wild-type. Thus, it is possible that a large part of the drying-induced genes during *Arabidopsis* seed maturation may function in an ABA-independent manner.

The effects of the drying regime extended beyond the transcriptional landscape. We observed global co-translational regulation of mRNAs by monitoring the CTRD pathway under SD. SD not only decreased overall CTRD activity but also caused a shift in the mRNA pool undergoing CTRD. This shift moved from a gene pool primarily associated with metabolism and development to one that is more related to stress (Fig. 6c and Fig. S16f). For a specific subset of maturation-related genes, SD resulted in increased read accumulation near the start codon, indicating higher ribosome stalling in these genes (Table S10). Additionally, our analysis revealed that SD treatment promotes a massive shutdown of translation and stabilization of *LEA* transcripts (Fig. 6). This coordinated mechanism contributes to seed maturation. Indeed, SD treatment induces an accumulation of 5’P reads 13 nt upstream of the start codon, corresponding to ribosomes stalled at the initiation step (with the ATG in the P-site). Interestingly, a similar profile is observed in dry seeds (Bai *et al*., 2026). It is tempting to propose that SD treatment facilitates the establishment of the translational program required for seed germination.

Overall, our study provides fundamental insights into the mechanisms by which seeds perceive and respond to drying, advancing our basic understanding of the molecular regulation of seed maturation and DT. These findings can directly inform post-harvest, storage, and post-priming drying practices to enhance seed quality and storability.

## Materials and Methods

### Plant material and growth conditions

Experiments were performed using the *Arabidopsis thaliana* (L.) Heynh. accession Columbia (Col-0) and the *aba2-1* mutant (Leon-Kloosterziel *et al*., 1996). Additionally, four transgenic lines expressing a LEA_4 protein fused with the Venus fluorescent protein under their native promoter (proLEA_4::LEA_4:Venus) were utilized for western blotting (Ginsawaeng *et al*., 2021). Plants were grown in 4x4 cm Rockwool blocks in a growth chamber at 20°C/18°C (day/night) under a 16-h photoperiod of artificial light (150 μmol m-2 s-1) and ∼70% relative humidity (RH). Plants were watered three times per week with the standard Hyponex nutrient solution (He *et al*., 2014). Once plants bolted, flowers were tagged just prior to anthesis, i.e., on the day of pollination. After tagging, the time was recorded as days after pollination (DAP).

The following T-DNA mutant lines were obtained from Nottingham Arabidopsis Stock Center (NASC). WRKY46 (SALK_134310C), MYC2 (SALK017005C), ERF13 (SALK_019879), NUDT7 (SALK_046441), XTH22 (SAIL_422_D11), NILR2 (SAIL_878_C01), CYP74A (SALK_017756), RHA2B (SALK_12-233). Single plants were genotyped with Phire Plant Direct PCR Kit (Thermo Scientific, F130WH). More details about T-DNA lines and primers can be found in Table S7.

Seeds from heterozygous transgenic *A. thaliana* lines (from T2 seeds) of the four LEA_4 seed-expressed proteins were kindly provided by Dr. Ellen Zuther (Max Planck Institute of Molecular Plant Physiology) (Ginsawaeng *et al*., 2021). These lines corresponded to the genes *LEA7* (*AT1G52690*), *LEA29* (*AT3G15670*), *LEA30* (*AT3G17520*), and *LEA48* (*AT5G44310*). These lines contained the LEA_4 gene under its native promoter fused with a fluorescent protein Venus tag (proLEA_4::LEA_4:Venus) and a kanamycin resistance gene in a Col-0 background. Homozygous lines were selected and used for further experiments.

### Drying regimes

Following harvest, seeds underwent four drying treatments prior to phenotyping and RNA extraction – Fresh, fast drying (FD), slow drying (SD), and slow drying followed by fast drying (SDFD). Fresh samples were prepared with seeds that were immediately removed from the siliques after harvest using fine tweezers with no further treatments. FD samples comprised seeds removed from the siliques after harvest and placed on filter papers (Sartorius, 50mm diameter, white, grade 3hw) pre-soaked with 300 μL of demineralized water (demi H_2_O). Subsequently, the seeds were transferred to a drying cabinet with 30% RH at 20 °C and dried for 2 days. For SD samples, intact siliques with the seeds were placed in a drying cabinet with 30% RH at 20 °C, and seeds were allowed to dry inside the silique for 2 days. SDFD samples combined 2 days of SD followed by 2 days of FD. Seeds were removed from the siliques following SD and prior to FD. For all phenotypic assays and RNA extraction related to samples treated under different drying regimes, seeds were harvested at 10,12, 14, 16, and 26 DAP.

### Water content and desiccation tolerance (DT)

Seed water content was determined using a gravimetric approach on a fresh weight basis. Water content during seed maturation was measured using 4 biological replicates; each replicate consisted of a pool of seeds from 5 siliques. Siliques were harvested at different time points starting from 10 DAP until 26 DAP with 2 days of interval. Seeds were extracted from the silique using fine tweezers, and the fresh weight was measured within 5 minutes of extraction. The dry weight was measured following 24 hours of drying at 105 °C. The water content was expressed as grams H_2_O per gram Dry weight (g/gDW) and relative water content (%). Water content of seeds following drying treatments was measured by taking the fresh weight after the drying treatments using 3 biological replicates, each replicate consisting of seeds pooled from 5 siliques. The dry weight was measured after 24 hours of drying at 105 °C.

To determine DT during maturation, seeds harvested at 10 – 26 DAP were germinated on two layers of blue blotter paper (Anchor Paper Co., St Paul, MN, USA) soaked in 40mL of demi H_2_O supplemented with 1mM of GA_4+7_ and 10mM of KNO_3_. For each time point, 4 biological replicates were used, with each replicate comprising seeds from 1-2 siliques. DT survival after drying treatments was measured by germinating untreated (Fresh), FD treated, and SDFD treated seeds on two layers of blue blotter paper soaked in 40mL of demi H_2_O supplemented with 1mM of GA_4+7_ and 10mM of KNO_3_. For each time point and treatment combination, 6 biological replicates were used, with each replicate comprising seeds from 1-2 siliques. No germination was performed with seeds treated with SD only, since the seeds were not fully desiccated.

### ABA measurement

ABA measurement was performed as described in Lamers *et al*., 2025. Three biological replicates were used, with each replicate consisting of 6 siliques (approximately 300 seeds). Seeds were dissected from the silique tissues without drying treatments (Fresh), or after two days of SD. For FD samples, seeds were dissected from siliques, and both dissected seeds and silique tissues were dried for two days in a drying cabinet (30%RH, 20 °C). All samples were frozen and ground to a fine powder using a 5/32cm stainless steel bead in a mixer mill (Retsch MM400). Ground material was extracted in 10% MEOH containing 100nM stable isotope-labeled internal standard. Further extraction and measurement were as described in Lamers *et al*., 2025.

### RNA-extraction and sequencing

Seeds from 6 siliques at different DAP and submitted to the different drying regimes were combined in a 2mL tube and frozen in liquid nitrogen. Seeds were ground to fine powder using a 5/32 stainless steel bead in a mixer mill (Retsch MM400). Total RNA was isolated as described by Maia *et al*., 2011. In brief, fine powder was dissolved in 800µL extraction buffer containing DTT and proteinase K. After centrifugation, RNA was precipitated overnight with Lithium Chloride. Precipitated RNA was DNase-treated. RNA poly-A enrichment library preparation and RNA sequencing were carried out by Novogene using the Illumina NovaSeq 6000 platform.

### RNA-seq analysis

Raw sequencing reads were processed using the nf-core/rnaseq pipeline (v3.9) (Ewels *et al*., 2020) executed via the Nextflow workflow manager (v22.10.6) (Di Tommaso *et al*., 2017). The pipeline was executed with default parameters. Initially, raw read quality was assessed using FastQC (v0.11.9), followed by trimming using Trim Galore (v0.6.7), and quality was re-evaluated with FastQC. Trimmed reads were aligned to the reference genome using STAR (v2.7.10a) (Dobin *et al*., 2013). Gene-level expression quantification was subsequently performed from the STAR alignments using Salmon (v1.5.2) (Patro *et al*., 2017). Finally, results and quality control metrics from all pipeline steps were aggregated and summarized into a single report using MultiQC (v1.13) (Ewels *et al*., 2016). The resulting salmon quant output files were used to load the raw read counts for each RNA-seq sample. The raw reads were used as input for differential gene expression analysis using DESeq2 (v1.46.0) (Love *et al*., 2014). For each pairwise comparison, significant genes were filtered based on a Log2FoldChange of > 1.5 or < -1.5 with an adjusted p-value less than 0.01. The filtered genes were considered differentially expressed genes (DEGs).

The fastq files corresponding to round 2 of the SeedMatExplorer (Artur *et al*., 2024) were downloaded from NCBI (PRJNA1129160) using the Sequence Read Archive (SRA) Toolkit (v3.0.3). The reads were mapped to the *A. thaliana* TAIR10 reference genome using the nf-core/rnaseq pipeline as mentioned above.

### Module detection and gene regulatory network (GRN) inference

Gene co-expression modules were determined using the WGCNA package (v1.73) in R using Transcript per million (TPM) counts (Langfelder & Horvath, 2008). The RNA-seq data of *A. thaliana* maturation time course from the “SeedMatExplorer” was included in the analysis (Artur *et al*., 2024). Lowly expressed genes were filtered out by taking genes that had at least a TPM > 2 in three samples. The modules were constructed using only Col-0 samples (mutants were excluded). A signed network was constructed using a power (soft threshold) of 16, minModuleSize to 30, and mergeCutHeight to 0.2. This resulted in 14 gene modules named from M1 to M14. A correlation between the modules and four traits – germination %, relative water content (RWC), ABA content in seeds, and siliques-was performed using the “cor” function within WGCNA. The germination percentage for maturation time points of freshly harvested seeds was obtained from the SeedMatExplorer (Artur *et al*., 2024) (Table S5).

Gene regulatory network (GRN) inference was performed for modules of interest using GRNBoost2 under the arboreto package (v.0.1.6) in Python (v3.9.13) (Moerman *et al*., 2019). A list of transcription factors was downloaded from the Plant Transcription Factor Database (PlantTFDB) (Jin *et al*., 2017). The TFs within each module were identified using this list. GRNBoost2 was executed using the subsetted expression data and TFs in each module. The top 1000 connections were used to build the GRN.

### Gene ontology (GO) enrichment

GO enrichment analysis was performed using the TopGO package (v2.58.0) in R (Adrian Alexa, 2023). The GoSlim file was downloaded from TAIR. Enrichment was performed on the target using all protein-coding genes as background. GO terms with an FDR value of < 0.001 and a log2FC of > 0.5 were selected. Similar GO terms were merged using the rrvgo package (v1.18.0) in R (Sayols, 2023).

### 5’P sequencing and analysis

5′Pseq library was prepared for samples of 12, 14, and 16 DAP Fresh and SD samples as described previously (Carpentier *et al*., 2024). Raw reads were trimmed to 50 pb before mapping. Metagene analysis was performed using FIVEPSEQ software v1.0.0 (Nersisyan *et al*., 2020). ‘meta_counts_START.txt’ and ‘meta_counts_STOP.txt’ files were used to analyze 5′P reads accumulation around start and stop positions. The translational termination stalling index (TSI) was defined as the ratio of the number of 5′P read ends at the ribosome boundary (16–17 nt upstream from stop codons) to the mean number of 5′P read ends within the flanking 100 nt. Transcripts with a TSI value higher than 2 in WT were used to assess CTRD activity in Fresh and SD samples of Col-0.The - 13nt peaks near the start codon were compared between the SD and Fresh samples by taking the read count at -13nt before the start codon in the SD sample and dividing it by the read count at -13nt before the start codon in the Fresh sample from the same time point. Genes with a ratio > 1 were retained. For these genes, the -13nt start codon peaks were further compared to the -16/-17nt stop codon peaks for each SD sample. The comparison was made by dividing the -13nt read count by the average read count of the -16 and -17nt before the stop codon for each SD sample. Again, genes with a ratio > 1 were retained.

### Western blotting

Seeds of 16 DAP were treated under different drying regimes – Fresh (no drying), FD, SD, and SDFD. Following the treatment, the seeds were frozen in liquid nitrogen before being transferred to a -80 °C freezer. Grinding of frozen samples was done by shaking twice for 45 seconds each time at 28Hz with the grinding machine. A volume of 300 μL Lysis buffer was added to each tube and placed on ice. The lysis buffer comprised 400 mM Tris-HCl (pH 8.0), 1.25% sucrose, 200 mM LiCl, 35 mM MgCl_2_, 5 mM DTT, 5 mM EGTA, and 1% LiDS. This was achieved by combining 2 mL of 2 M Tris-HCl, 1.25 mL of 10% sucrose, 0.25 mL of 8 M LiCl, 0.35 mL of MgCl_2_ stock, 0.4 mL of 125 mM DTT, 0.1 mL of 0.5 M EGTA, and 100 µL of 10% LiDS. One mini tablet of protease inhibitor cocktail was dissolved into the mixture, and the solution was subsequently brought to a final volume of 10 mL using Milli-Q (MQ) water. Following incubation in the lysis buffer, the tubes were thoroughly vortexed, incubated on ice for 10 minutes, and then centrifuged at maximum speed (≥13,000 × g) for 20 minutes at 4 °C to separate the pellet from the supernatant. The supernatant was transferred into fresh 1.5 mL Eppendorf tubes. The total protein concentration in each sample was determined using the Bradford assay (Bradford reagent, Bio-Rad). For all samples, 5x Loading dye was added to all supernatants in a ratio of 1 part buffer + 4 parts supernatant. Samples were heated at 95 °C for 5 minutes and stored at -80 °C for later use. Samples were preheated at 95 °C for 5 minutes. A total of 15µg protein was loaded onto the gel. A volume of 5 μL precision plus protein dual color marker (Bio-Rad) was loaded as a marker, eGFP was loaded as a positive control, and dry Col-0 seeds were loaded as a negative control. Electrophoresis for stacking gel was run for 15 minutes at 50 volts, and then at 100 volts for approximately 120 minutes for separating gel.

The gel was incubated in the transfer buffer for 5 minutes, then transferred to a nitrocellulose membrane (Bio-Rad) with bottom and top cushions to form a transfer sandwich. The sandwich was transferred to the Turbo Blotter drawer (Bio-Rad). Blotting was done by selecting the ‘Turbo transfer’, followed by ‘Mixed MW preset program (25 V, 1.0 A, 7 minutes) settings, and finally selecting the loading cassette to start the transfer. The blot was blocked with 5 mL of a solution of 5% fat free milk powder in TBS-T (20 mM Tris-HCl, 150 mM NaCl, 0.1% Tween-20, pH 7.6), using a roller for 2 hours at room temperature. After blocking, the blot was washed for 10 minutes in TBS-T. The blot was transferred to a 50 mL tube containing 5 mL 3% fat free milk in TBS-T solution and 5 μL (1:1000 dilution) anti-GFP 3H9 monoclonal primary antibody (Chromotek). The 50 mL tube containing the blot was incubated overnight on a roller at 4°C. The blot was washed 3 times with TBS-T and then incubated for 1 hour with a solution containing 5 mL 3% fat free milk in TBS-T and 1 μL (1:5000) Anti-Rat IgG-HRP secondary antibody (Agrisera). Afterward, the blots were washed five times in TBS-T. The blots were developed for imaging using 350 μL each of the two Enhanced Chemiluminescence (ECL) substrates.

Images of the blots were taken with the ChemiDoc imaging machine using colorimetric and chemiluminescence settings to image the marker and the protein, respectively.

## Data availability

The RNAseq and 5’Pseq data have been deposited to the National Center for Biotechnology Information (NCBI) under the Bioproject ID PRJNA1444691.

## Acknowledgements

We thank Dr. Helen Zuther and Dr. Orarat Ginsawaeng (Max Planck Institute of Molecular Plant Physiology) for providing the LEA:Venus lines. Julia Damen and Hylke Schakel for their contribution with selection of LEA_4 mutant lines. Lars Bakermans, Kees Ketting, Sjors van der Horst and Anne-Elodie Receveur for their input on the degradome-seq data analyses. Francel Verstappen for the contribution with hormone measurements and all colleagues from the Wageningen Seed Science Centre and the Laboratory of Plant Physiology from Wageningen University & Research for their help with experimentation.

This work was funded by The Dutch Research Council (NWO) ENW Veni (project Fine Drying VI.Veni.202.038) to M.A.S.A., NWO VICI (project Seeds4Ever 17047) to L.B., and the French-Netherlands mobility program Campus France-NUFFIC Van Gogh (2022–2023), with support from the French Ministry for Europe and Foreign Affairs (MEAE), the French Ministry of Higher Education and Research (MESR), and the Netherlands Universities Foundation for International Cooperation (NUFFIC) to R. M., M-C.C and L. B..

## Supplementary figure legends

**Fig. S1**. Changes in water content (gDW) and dry weight (g) in seeds of *Arabidopsis thaliana* during maturation. Red and green lines indicate the dry weight (g) and water content (gDW), respectively. The grey area around the line indicates confidence interval. DS indicates fully matured dry seeds.

**Fig. S2**. ABA dynamics in maturing seeds of *Arabidopsis thaliana* in response to fast and slow drying. ABA was extracted either from seed or silique tissue of a total of six siliques containing approximately 300 seeds. (a) ABA levels in seed tissue (∼300 seeds). (**b**) ABA levels in silique tissue from six siliques. DS indicates the ABA levels in mature dry seeds (**a**) or mature dry siliques (**b**). Asterisks indicates significant difference of FD and SD samples compared to Fresh seeds of the same time point using the Wilcoxon-rank sum test. Error bars indicate standard deviation calculated from three replicates (n=3).

**Fig. S3**. Hierarchical clustering of the RNA-seq samples based on distance. Colored boxes indicate the two major clusters. Fresh indicates untreated seeds, and FD, SD, and SDFD represent the three different drying regimes. 26 DAP seeds represent mature dry seeds (DS).

**Fig. S4**. (**a**) Number of differentially expressed genes (DEGs) that are up- or downregulated in SDFD samples compared to Fresh samples for each time point. Counts include genes that have a significance of p-adj < 0.01. (**b**) Venn diagram showing core up (left) or downregulated (right) genes at all time points after SDFD treatment.

**Fig. S5**. (**a**) Upset plot showing the overlap between SD core downregulated genes and FD downregulated genes from different time points. (**b**) Upset plot showing the overlap between SD core upregulated genes and FD upregulated genes at different time points. The bars indicate an inclusive intersection between pairs.

**Fig. S6**. PCA showing clustering of RNA-seq samples maturing seeds generated in this study (10-16 DAP) and the SeedMatExplorer (Artur et al., 2024, 12-26 DAP). From the present study, only RNA-seq samples of freshly harvested seeds without any further drying treatments were shown. Transcripts per million (TPM) count of all genes was used to calculate PC1 and PC2.

**Fig. S7**. Heatmap showing the top 1ten GO terms enriched in each gene module. Color intensity indicates -log2 (FDR). Red-colored text indicates GO terms shared across multiple modules.

**Fig. S8**. Gene expression dynamics of GRN of M5 and M6 modules in response to FD. (**a**) Z-score changes of genes in the M5 module in response to FD. (**b**) Z-score changes of genes in the M6 module in response to FD. Nodes corresponding to major transcription factors (TFs) are shown in each network.

**Fig. S9**. Transcriptional profile of eight genes showing contrasting patterns under FD and SD. Error bars indicate standard error calculated from three replicates (n=3).

**Fig. S10**. Barplot showing the longevity (p50) of the seeds of different T-DNA lines following a controlled deterioration test (CDT) at 38°C and 75% RH. The mutant genes represent the six genes that overlap between FD up and SD/SDFD down, i.e., genes with contrasting responses to the drying regimes. The wild type (Col-0) was used as a control. Genotype effects on the p50 were determined using the Kruskal-Wallis test followed by Dunn’s test with multiple corrections. Significant differences are indicated using letters. Error bars indicate standard error calculated from six replicates (n=6).

**Fig. S11**. Maximum germination percentage (gMax, %) after 0, 7, 14, 21, 35, and 42 days of controlled deterioration test (CDT) at 38°C and 75% RH. Genotype effects on both gMax were determined using the Kruskal-Wallis test followed by Dunn’s test with multiple corrections. Significant differences are indicated using letters. Error bars indicate standard error calculated from six replicates (n=6) over the CDT time points.

**Fig. S12**. GO enrichment analysis of genes overlapping between Col-0 core SD downregulated and *aba2-1* 14 DAP SD downregulated genes. Size of the bubbles indicate -log2(FDR) values for each GO term.

**Fig. S13**. Gene expression level in transcripts per million (TPM) of *ABI3* and *ABI5* genes based on the RNA-seq data. (**a**) Expression of *ABI3* and *ABI5* genes from 10 - 16 DAP in wild type (Col-0). (**b**) Expression of *ABI3* and *ABI5* genes at 14 DAP in the *aba2-1* mutant. Error bars indicate standard error calculated from three replicates (n=3).

**Fig. S14**. The Fast Fourier transformation (FFT) signal near the (**a**) start and (**b**) stop codon regions. Colors indicate the replicates (n=3).

**Fig. S15**. (**a**) Metagene profile near the START codon of genes (n = 862) with a –13 nt read count ratio > 1 between SD and Fresh samples (read count at –13 nt in SD sample/ read count at –13 nt in Fresh) at 12, 14, and 16 DAP. (**b**) Top 25 GO terms enriched in these genes. Color indicates the -log10 (FDR) value, and size indicates the number of significant genes in each GO category.

**Fig. S16**. Comparison of genes undergoing CTRD (TSI >= 2) between Fresh and SD samples. (**a**) Upset plot showing the overlap in CTRD genes between different timepoint-treatment combinations. Bars highlighted in red, blue, and yellow indicate CTRD genes shared between all, Fresh, and SD samples, respectively. Gene counts are inclusive. (**b**) Venn diagram showing overlap between genes undergoing CTRD in Fresh and SD samples. (**c**) Distribution of Termination Stalled Index (TSI) of CTRD genes unique to Fresh (left) and SD (right) samples over different timepoint-treatment combinations. (**d**) Distribution of Termination Stalled Index (TSI) of CTRD genes unique to SD. (**e**) Distribution of Termination Stalled Index (TSI) of CTRD genes shared between Fresh and SD. n = 3, biological replicates for all samples except 12 DAP Fresh, where n = 2. Statistical significance was determined using the Wilcoxon rank sum test. (**f**) GO terms enriched in CTRD genes that are specific to Fresh or SD or shared between both.

**Fig. S17**. Metagene profiles around start (left) and stop codons (right) of gene undergoing CTRD (TSI>2). (**a**) Metagene profiles of CTRD genes unique to Fresh samples. (**b**) Metagene profiles of CTRD genes unique to SD samples. (**c**) Metagene profiles of CTRD genes shared between Fresh and SD samples.

## Supplementary table legends

**Table S1**. Lists of core up and downregulated genes under FD, SD, and SDFD.

**Table S2**. List of genes with their categories based on overlap of core FD, SD, and SDFD up- and downregulated DEGs.

**Table S3**. Full list of *Arabidopsis* genes along with their gene modules identified using WGCNA.

**Table S4**. Full list GO terms enriched in each gene module from M1 to M14.

**Table S5**. Physiological traits associated with RNA-seq samples.

**Table S6**. Top 1,000 connections in the M5 and M6 GRN built with GRNBoost2.

**Table S7**. List of FD-activated genes whose T-DNA mutants were used in the controlled deterioration test (CDT).

**Table S8**. List of up- and downregulated DEGs in 14 DAP FD and SD seeds of *aba2-1* mutant.

**Table S9**. List of genes with a relative read count ratio > 1 at -13nt peak ratio between SD and Fresh.

**Table S10**. List of genes with a relative read count ratio > 1 between –13nt region around START codon compared to the average of -16/-17nt around the STOP codon in SD samples.

## Notes

### Competing Interest Statement

The authors have declared no competing interest.

## References

Adams CA, Fjerstad MC, Rinne RW. 1983. Characteristics of Soybean Seed Maturation: Necessity for Slow Dehydration. Crop Science 23: 265–267.

Adrian Alexa JR. 2023. topGO: Enrichment Analysis for Gene Ontology.

Angelovici R, Galili G, Fernie AR, Fait A. 2010a. Seed desiccation: a bridge between maturation and germination. Trends in Plant Science 15: 211–218.

Angelovici R, Galili G, Fernie AR, Fait A. 2010b. Seed desiccation: a bridge between maturation and germination. Trends in Plant Science 15: 211–218.

Artur MAS, Koetsier RA, Willems LAJ, Bakermans LL, Van Driel AD, Dongus JA, Dekkers BJW, Marques ACSS, Sami AA, Nijveen H, et al. 2024. SeedMatExplorer: The transcriptome atlas of Arabidopsis seed maturation.

Bai B, Qi R, Song W, Nijveen H, Bentsink L. 2026. Translational landscape during seed germination revealed by ribosome profiling. The Plant Journal 125: e70663.

Bai B, Schiffthaler B, Van Der Horst S, Willems L, Vergara A, Karlström J, Mähler N, Delhomme N, Bentsink L, Hanson J. 2023. SeedTransNet: a directional translational network revealing regulatory patterns during seed maturation and germination (P Manavella, Ed.). Journal of Experimental Botany 74: 2416–2432.

Bai B, Van Der Horst S, Cordewener JHG, America TAHP, Hanson J, Bentsink L. 2020. Seed-Stored mRNAs that Are Specifically Associated to Monosomes Are Translationally Regulated during Germination. Plant Physiology 182: 378–392.

Baud S, Boutin J-P, Miquel M, Lepiniec L, Rochat C. 2002. An integrated overview of seed development in Arabidopsis thaliana ecotype WS. Plant Physiology and Biochemistry 40: 151–160.

Bies N, Aspart L, Carles C, Gallois P, Delseny M. 1998. Accumulation and degradation of Em proteins in Arabidopsis thaliana: evidence for post-transcriptional controls. Journal of Experimental Botany 49: 1925–1933.

Blackman SA, Obendorf RL, Leopold AC. 1992. Maturation Proteins and Sugars in Desiccation Tolerance of Developing Soybean Seeds. Plant Physiology 100: 225–230.

Butler LH, Hay FR, Ellis RH, Smith RD, Murray TB. 2009. Priming and re-drying improve the survival of mature seeds of Digitalis purpurea during storage. Annals of Botany 103: 1261–1270.

Carpentier M-C, Deragon J-M, Jean V, Be SHV, Bousquet-Antonelli C, Merret R. 2020. Monitoring of XRN4 Targets Reveals the Importance of Cotranslational Decay during Arabidopsis Development. Plant Physiology 184: 1251–1262.

Carpentier M-C, Receveur A-E, Boubegtitene A, Cadoudal A, Bousquet-Antonelli C, Merret R. 2024. Genome-wide analysis of mRNA decay in Arabidopsis shoot and root reveals the importance of co-translational mRNA decay in the general mRNA turnover. Nucleic Acids Research 52: 7910–7924.

Chatelain E, Hundertmark M, Leprince O, Gall SL, Satour P, Deligny-Penninck S, Rogniaux H, Buitink J. 2012. Temporal profiling of the heat-stable proteome during late maturation of Medicago truncatula seeds identifies a restricted subset of late embryogenesis abundant proteins associated with longevity. Plant, Cell & Environment 35: 1440–1455.

Cheng ZJ, Zhao XY, Shao XX, Wang F, Zhou C, Liu YG, Zhang Y, Zhang XS. 2014. Abscisic Acid Regulates Early Seed Development in Arabidopsis by ABI5-Mediated Transcription of SHORT HYPOCOTYL UNDER BLUE1. The Plant Cell 26: 1053–1068.

Corbineau F, Picard MA, Fougereux J-A, Ladonne F, Côme D. 2000. Effects of dehydration conditions on desiccation tolerance of developing pea seeds as related to oligosaccharide content and cell membrane properties. Seed Science Research 10: 329–339.

Dannfald A, Carpentier M-C, Merret R, Favory J-J, Deragon J-M. 2025. Plant response to intermittent heat stress involves modulation of mRNA translation efficiency. Plant Physiology 197: kiae648.

Deragon J-M, Merret R. 2025. Co-translational mRNA decay in plants: recent advances and future directions (C Merchante, Ed.). Journal of Experimental Botany: eraf146.

Di Tommaso P, Chatzou M, Floden EW, Barja PP, Palumbo E, Notredame C. 2017. Nextflow enables reproducible computational workflows. Nature Biotechnology 35: 316–319.

Diamond J. 2002. Evolution, consequences and future of plant and animal domestication. Nature 418: 700–707.

Dobin A, Davis CA, Schlesinger F, Drenkow J, Zaleski C, Jha S, Batut P, Chaisson M, Gingeras TR. 2013. STAR: ultrafast universal RNA-seq aligner. Bioinformatics 29: 15–21.

Dombrecht B, Xue GP, Sprague SJ, Kirkegaard JA, Ross JJ, Reid JB, Fitt GP, Sewelam N, Schenk PM, Manners JM, et al. 2007. MYC2 Differentially Modulates Diverse Jasmonate-Dependent Functions in Arabidopsis. The Plant Cell 19: 2225–2245.

Dong N-Q, Lin H-X. 2021. Contribution of phenylpropanoid metabolism to plant development and plant-environment interactions. Journal of Integrative Plant Biology 63: 180–209.

Ewels P, Magnusson M, Lundin S, Käller M. 2016. MultiQC: summarize analysis results for multiple tools and samples in a single report. Bioinformatics 32: 3047–3048.

Ewels PA, Peltzer A, Fillinger S, Patel H, Alneberg J, Wilm A, Garcia MU, Di Tommaso P, Nahnsen S. 2020. The nf-core framework for community-curated bioinformatics pipelines. Nature Biotechnology 38: 276–278.

Fàbregas N, Yoshida T, Fernie AR. 2020. Role of Raf-like kinases in SnRK2 activation and osmotic stress response in plants. Nature Communications 11: 6184.

Fabrissin I, Sano N, Seo M, North HM. 2021. Ageing beautifully: can the benefits of seed priming be separated from a reduced lifespan trade-off? (S Penfield, Ed.). Journal of Experimental Botany 72: 2312–2333.

Fonseca S, Chini A, Hamberg M, Adie B, Porzel A, Kramell R, Miersch O, Wasternack C, Solano R. 2009. (+)-7-iso-Jasmonoyl-L-isoleucine is the endogenous bioactive jasmonate. Nature Chemical Biology 5: 344–350.

Gazzarrini S, Song L. 2024. LAFL Factors in Seed Development and Phase Transitions. Annual Review of Plant Biology 75: 459–488.

Ginsawaeng O, Heise C, Sangwan R, Karcher D, Hernández-Sánchez IE, Sampathkumar A, Zuther E. 2021. Subcellular Localization of Seed-Expressed LEA_4 Proteins Reveals Liquid-Liquid Phase Separation for LEA9 and for LEA48 Homo- and LEA42-LEA48 Heterodimers. Biomolecules 11: 1770.

González-Morales SI, Chávez-Montes RA, Hayano-Kanashiro C, Alejo-Jacuinde G, Rico-Cambron TY, De Folter S, Herrera-Estrella L. 2016. Regulatory network analysis reveals novel regulators of seed desiccation tolerance in Arabidopsis thaliana. Proceedings of the National Academy of Sciences 113.

Guo Y, Chen Y, Wang Y, Wu X, Zhang X, Mao W, Yu H, Guo K, Xu J, Ma L, et al. 2023a. The translational landscape of bread wheat during grain development. The Plant Cell 35: 1848–1867.

Guo R, Yu X, Gregory BD. 2023b. The identification of conserved sequence features of co-translationally decayed mRNAs and upstream open reading frames in angiosperm transcriptomes. Plant Direct 7: e479.

Hasanuzzaman M, Nahar K, Anee TI, Fujita M. 2017. Glutathione in plants: biosynthesis and physiological role in environmental stress tolerance. Physiology and Molecular Biology of Plants 23: 249–268.

Hay F. 1995. Seed Maturity and the Effects of Different Drying Conditions on Desiccation Tolerance and Seed Longevity in Foxglove (Digitalis purpurea L.). Annals of Botany 76: 639–647.

He H, De Souza Vidigal D, Snoek LB, Schnabel S, Nijveen H, Hilhorst H, Bentsink L. 2014. Interaction between parental environment and genotype affects plant and seed performance in Arabidopsis. Journal of Experimental Botany 65: 6603–6615.

Hoekstra FA, Golovina EA, Buitink J. 2001. Mechanisms of plant desiccation tolerance. Trends in Plant Science 6: 431–438.

Jin J, Tian F, Yang D-C, Meng Y-Q, Kong L, Luo J, Gao G. 2017. PlantTFDB 4.0: toward a central hub for transcription factors and regulatory interactions in plants. Nucleic Acids Research 45: D1040– D1045.

Jing Y, Lang S, Wang D, Xue H, Wang X-F. 2018. Functional characterization of galactinol synthase and raffinose synthase in desiccation tolerance acquisition in developing Arabidopsis seeds. Journal of Plant Physiology 230: 109–121.

Kermode AR. 1990. Regulatory mechanisms involved in the transition from seed development to germination. Critical Reviews in Plant Sciences 9: 155–195.

Lamers J, Zhang Y, Van Zelm E, Leong CK, Meyer AJ, De Zeeuw T, Verstappen F, Veen M, Deolu-Ajayi AO, Gommers CMM, et al. 2025. Abscisic acid signaling gates salt-induced responses of plant roots. Proceedings of the National Academy of Sciences 122: e2406373122.

Langfelder P, Horvath S. 2008. WGCNA: an R package for weighted correlation network analysis. BMC Bioinformatics 9: 559.

Leon-Kloosterziel KM, Van De Bunt GA, Zeevaart Jad, Koornneef M. 1996. Arabidopsis Mutants with a Reduced Seed Dormancy. Plant Physiology 110: 233–240.

Leprince O, Pellizzaro A, Berriri S, Buitink J. 2016. Late seed maturation: drying without dying. Journal of Experimental Botany: erw363.

Love MI, Huber W, Anders S. 2014. Moderated estimation of fold change and dispersion for RNA-seq data with DESeq2. Genome Biology 15: 550.

Maia J, Dekkers BJW, Provart NJ, Ligterink W, Hilhorst HWM. 2011. The Re-Establishment of Desiccation Tolerance in Germinated Arabidopsis thaliana Seeds and Its Associated Transcriptome (MA Blazquez, Ed.). PLoS ONE 6: e29123.

Marks RA, Ekwealor JTB, Artur MAS, Bondi L, Boothby TC, Carmo OMS, Centeno DC, Coe KK, Dace HJW, Field S, et al. 2025. Life on the dry side: a roadmap to understanding desiccation tolerance and accelerating translational applications. Nature Communications 16: 3284.

Meyer RS, Purugganan MD. 2013. Evolution of crop species: genetics of domestication and diversification. Nature Reviews Genetics 14: 840–852.

Moerman T, Aibar Santos S, Bravo González-Blas C, Simm J, Moreau Y, Aerts J, Aerts S. 2019. GRNBoost2 and Arboreto: efficient and scalable inference of gene regulatory networks (J Kelso, Ed.). Bioinformatics 35: 2159–2161.

Nersisyan L, Ropat M, Pelechano V. 2020. Improved computational analysis of ribosome dynamics from 5′P degradome data using fivepseq. NAR Genomics and Bioinformatics 2: lqaa099.

Oliver MJ, Farrant JM, Hilhorst HWM, Mundree S, Williams B, Bewley JD. 2020. Desiccation Tolerance: Avoiding Cellular Damage During Drying and Rehydration. Annual Review of Plant Biology 71: 435–460.

Ooms Jjj, Leon-Kloosterziel KM, Bartels D, Koornneef M, Karssen CM. 1993. Acquisition of Desiccation Tolerance and Longevity in Seeds of Arabidopsis thaliana (A Comparative Study Using Abscisic Acid-Insensitive abi3 Mutants). Plant Physiology 102: 1185–1191.

Patro R, Duggal G, Love MI, Irizarry RA, Kingsford C. 2017. Salmon provides fast and bias-aware quantification of transcript expression. Nature Methods 14: 417–419.

Pelechano V, Wei W, Steinmetz LM. 2015. Widespread Co-translational RNA Decay Reveals Ribosome Dynamics. Cell 161: 1400–1412.

Peredo EL, Cardon ZG. 2020. Shared up-regulation and contrasting down-regulation of gene expression distinguish desiccation-tolerant from intolerant green algae. Proceedings of the National Academy of Sciences 117: 17438–17445.

Prall W, Sharma B, Gregory BD. 2019. Transcription Is Just the Beginning of Gene Expression Regulation: The Functional Significance of RNA-Binding Proteins to Post-transcriptional Processes in Plants. Plant and Cell Physiology 60: 1939–1952.

Righetti K, Vu JL, Pelletier S, Vu BL, Glaab E, Lalanne D, Pasha A, Patel RV, Provart NJ, Verdier J, et al. 2015. Inference of Longevity-Related Genes from a Robust Coexpression Network of Seed Maturation Identifies Regulators Linking Seed Storability to Biotic Defense-Related Pathways. The Plant Cell: tpc.15.00632.

Roychoudhury A, Paul S, Basu S. 2013. Cross-talk between abscisic acid-dependent and abscisic acid-independent pathways during abiotic stress. Plant Cell Reports 32: 985–1006.

Sajeev N, Bai B, Bentsink L. 2019. Seeds: A Unique System to Study Translational Regulation. Trends in Plant Science 24: 487–495.

Sajeev N, Baral A, America AHP, Willems LAJ, Merret R, Bentsink L. 2022. The mRNA-binding proteome of a critical phase transition during Arabidopsis seed germination. New Phytologist 233: 251–264.

Samarah NH. 2006. Effect of air-drying immature seeds in harvested pods on seed quality of common vetch (Vicia sativa L.). New Zealand Journal of Agricultural Research 49: 331–339.

Samarah NH, Al-Mahasneh MM, Ghosheh HZ, Alqudah AM, Turk M. 2010. The influence of drying methods on the acquisition of seed desiccation tolerance and the maintenance of vigour in wheat (Triticum durum). Seed Science and Technology 38: 193–208.

Sanhewe AJ, Ellis RH. 1996. Seed development and maturation in Phaseolus vulgaris I. Ability to germinate and to tolerate desiccation. Journal of Experimental Botany 47: 949–958.

Sayols S. 2023. rrvgo: a Bioconductor package for interpreting lists of Gene Ontology terms. microPublication Biology 2023.

Shinozaki K, Yamaguchi-Shinozaki K. 2000. Molecular responses to dehydration and low temperature: differences and cross-talk between two stress signaling pathways. Current Opinion in Plant Biology 3: 217–223.

Sinnecker P, Braga N, Macchione ELA, Lanfer-Marquez UM. 2005. Mechanism of soybean (Glycine max L. Merrill) degreening related to maturity stage and postharvest drying temperature. Postharvest Biology and Technology 38: 269–279.

Sun P, Huang Y, Yang X, Liao A, Wu J. 2022. The role of indole derivative in the growth of plants: A review. Frontiers in Plant Science 13: 1120613.

Takahashi Y, Zhang J, Hsu P-K, Ceciliato PHO, Zhang L, Dubeaux G, Munemasa S, Ge C, Zhao Y, Hauser F, et al. 2020. MAP3Kinase-dependent SnRK2-kinase activation is required for abscisic acid signal transduction and rapid osmotic stress response. Nature Communications 11: 12.

Verdier J, Lalanne D, Pelletier S, Torres-Jerez I, Righetti K, Bandyopadhyay K, Leprince O, Chatelain E, Vu BL, Gouzy J, et al. 2013. A Regulatory Network-Based Approach Dissects Late Maturation Processes Related to the Acquisition of Desiccation Tolerance and Longevity of Medicago truncatula Seeds. PLANT PHYSIOLOGY 163: 757–774.

Veser J, Van Der Tuin J, Kodde J, Groot SPC, Van Der Sman RGM, Schutyser MAI. 2025. Effect of drying conditions on quality of primed cabbage seeds and ethanol degradation as promising quality parameter. Drying Technology 43: 753–770.

Wang C, Lyu Y, Zhang Q, Guo H, Chen D, Chen X. 2023. Disruption of BG14 results in enhanced callose deposition in developing seeds and decreases seed longevity and seed dormancy in Arabidopsis. The Plant Journal 113: 1080–1094.

Wang H, Li S, Li Y, Xu Y, Wang Y, Zhang R, Sun W, Chen Q, Wang X, Li C, et al. 2019. MED25 connects enhancer–promoter looping and MYC2-dependent activation of jasmonate signalling. Nature Plants 5: 616–625.

Yoshida T, Mogami J, Yamaguchi-Shinozaki K. 2014. ABA-dependent and ABA-independent signaling in response to osmotic stress in plants. Current Opinion in Plant Biology 21: 133–139.

Zander M, Lewsey MG, Clark NM, Yin L, Bartlett A, Saldierna Guzmán JP, Hann E, Langford AE, Jow B, Wise A, et al. 2020. Integrated multi-omics framework of the plant response to jasmonic acid. Nature Plants 6: 290–302.

Zhang Y, Xu P, Xue W, Zhu W, Yu X. 2023. Diurnal gene oscillations modulated by RNA metabolism in tomato. The Plant Journal 116: 728–743

