## Supplementary Figures for "Drying kinetics govern transcriptional and post-transcriptional reprogramming during seed maturation"

*Mariana A. S. Artur

**This PDF file includes:**

Figures S1 to 17

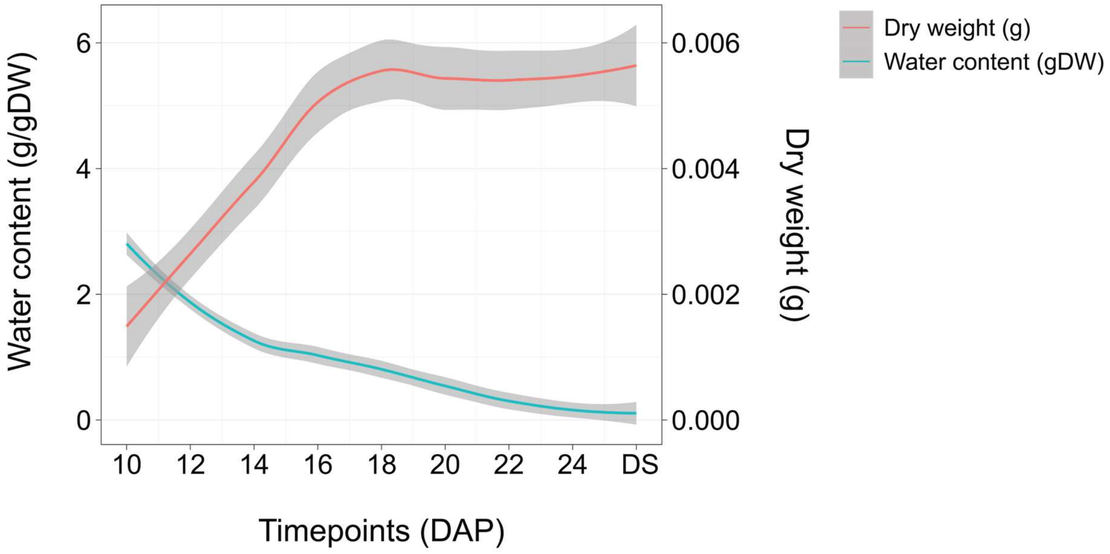


**Fig. S1**. Changes in water content (gDW) and dry weight (g) in seeds of *Arabidopsis thaliana* during maturation. Red and green lines indicate the dry weight (g) and water content (gDW), respectively. The grey area around the line indicates confidence interval. DS indicates fully matured dry seeds.

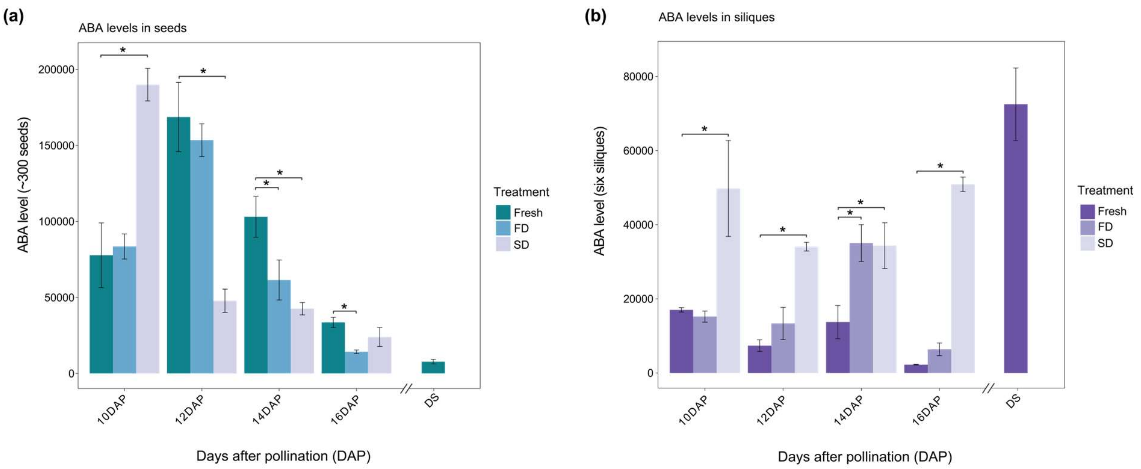


**Fig. S2**. ABA dynamics in maturing seeds of *Arabidopsis thaliana* in response to fast and slow drying. ABA was extracted either from seed or silique tissue of a total of six siliques containing approximately 300 seeds. **(a)** ABA levels in seed tissue (~300 seeds). **(b)** ABA levels in silique tissue from six siliques. DS indicates the ABA levels in mature dry seeds **(a)** or mature dry siliques **(b).** Asterisks indicates significant difference of FD and SD samples compared to Fresh seeds of the same time point using the Wilcoxon-rank sum test. Error bars indicate standard deviation calculated from three replicates (n=3).

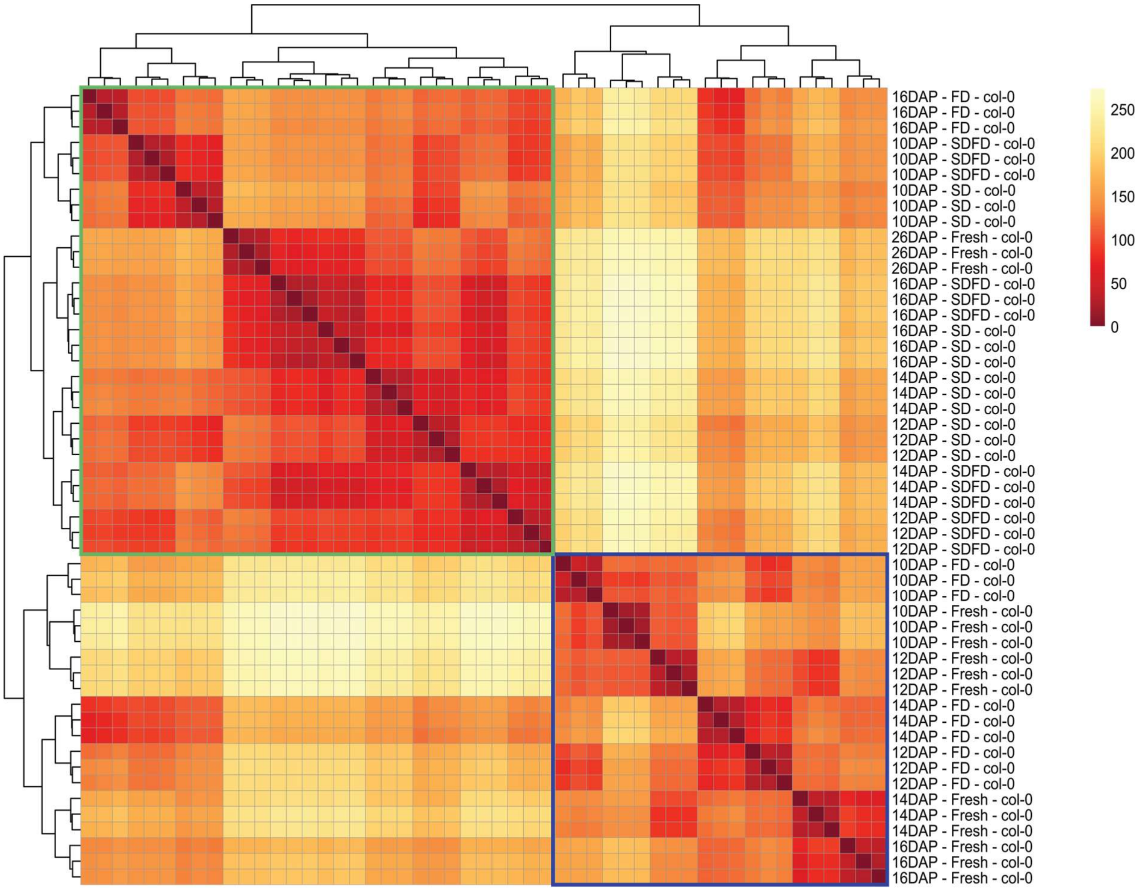


**Fig. S3**. Hierarchical clustering of the RNA-seq samples based on distance. Colored boxes indicate the two major clusters. Fresh indicates untreated seeds, and FD, SD, and SDFD represent the three different drying regimes. 26 DAP seeds represent mature dry seeds (DS).

**(a)**

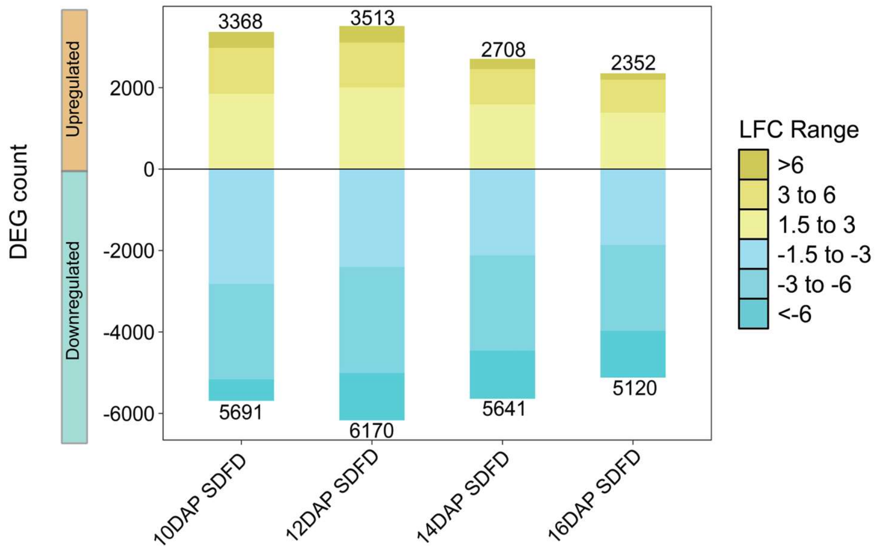


**(b)**

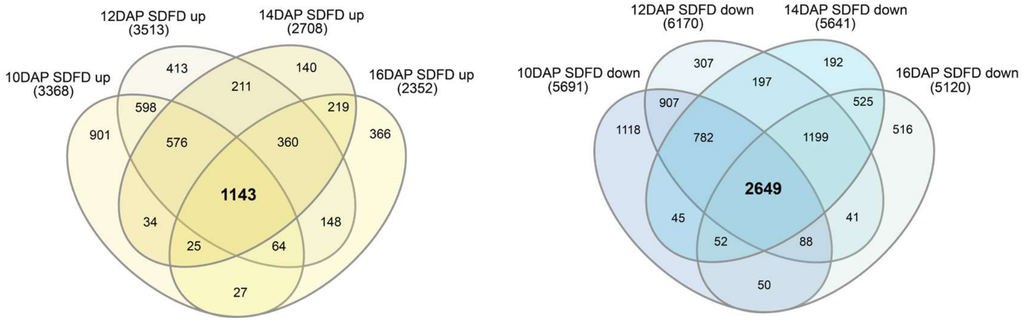


**Fig. S4. (a)** Number of differentially expressed genes (DEGs) that are up- or downregulated in SDFD samples compared to Fresh samples for each time point. Counts include genes that have a significance of p-adj < 0.01. (**b**) Venn diagram showing core up (left) or downregulated (right) genes at all time points after SDFD treatment.

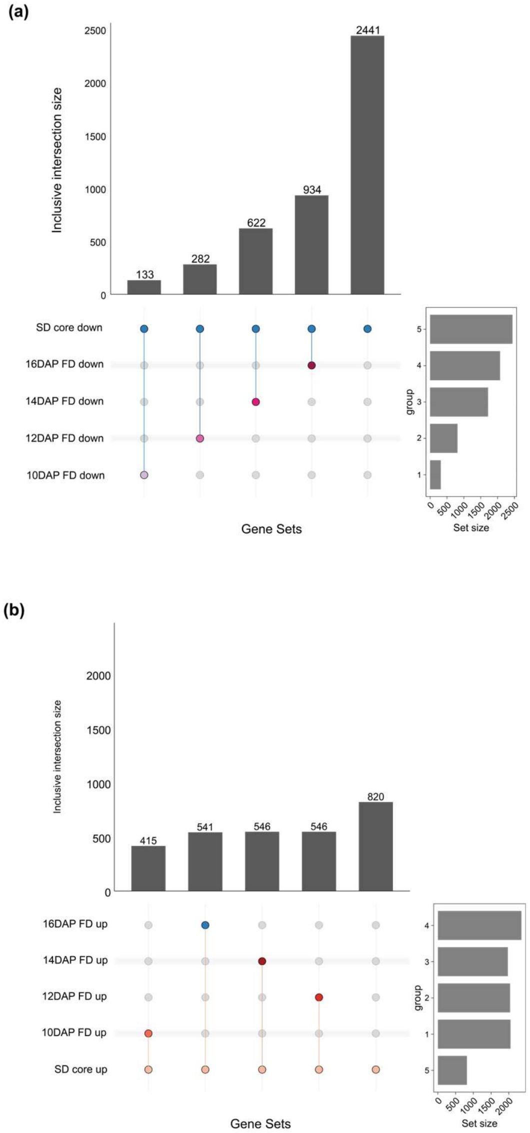


**Fig. S5. (a)** Upset plot showing the overlap between SD core downregulated genes and FD downregulated genes from different time points. **(b)** Upset plot showing the overlap between SD core upregulated genes and FD upregulated genes at different time points. The bars indicate an inclusive intersection between pairs.

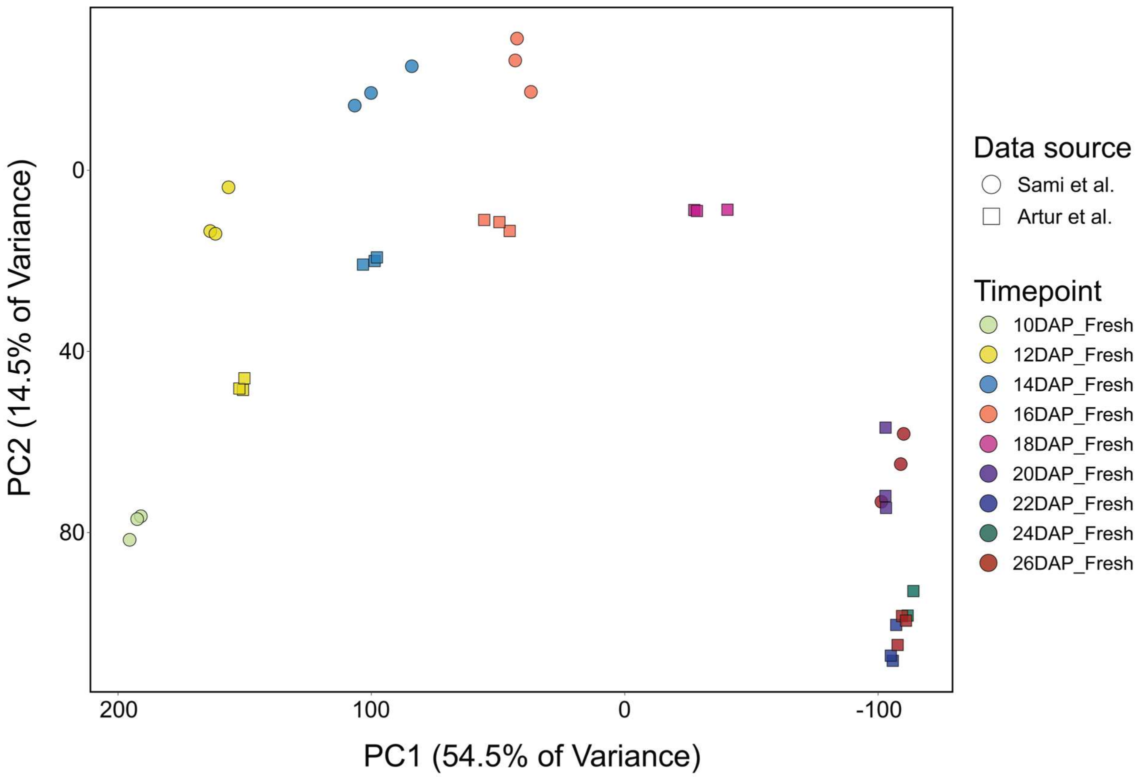


**Fig. S6**. PCA showing clustering of RNA-seq samples maturing seeds generated in this study (10-16 DAP) and the SeedMatExplorer (Artur et al., 2024, 12-26 DAP). From the present study, only RNA-seq samples of freshly harvested seeds without any further drying treatments were shown. Transcripts per million (TPM) count of all genes was used to calculate PC1 and PC2.

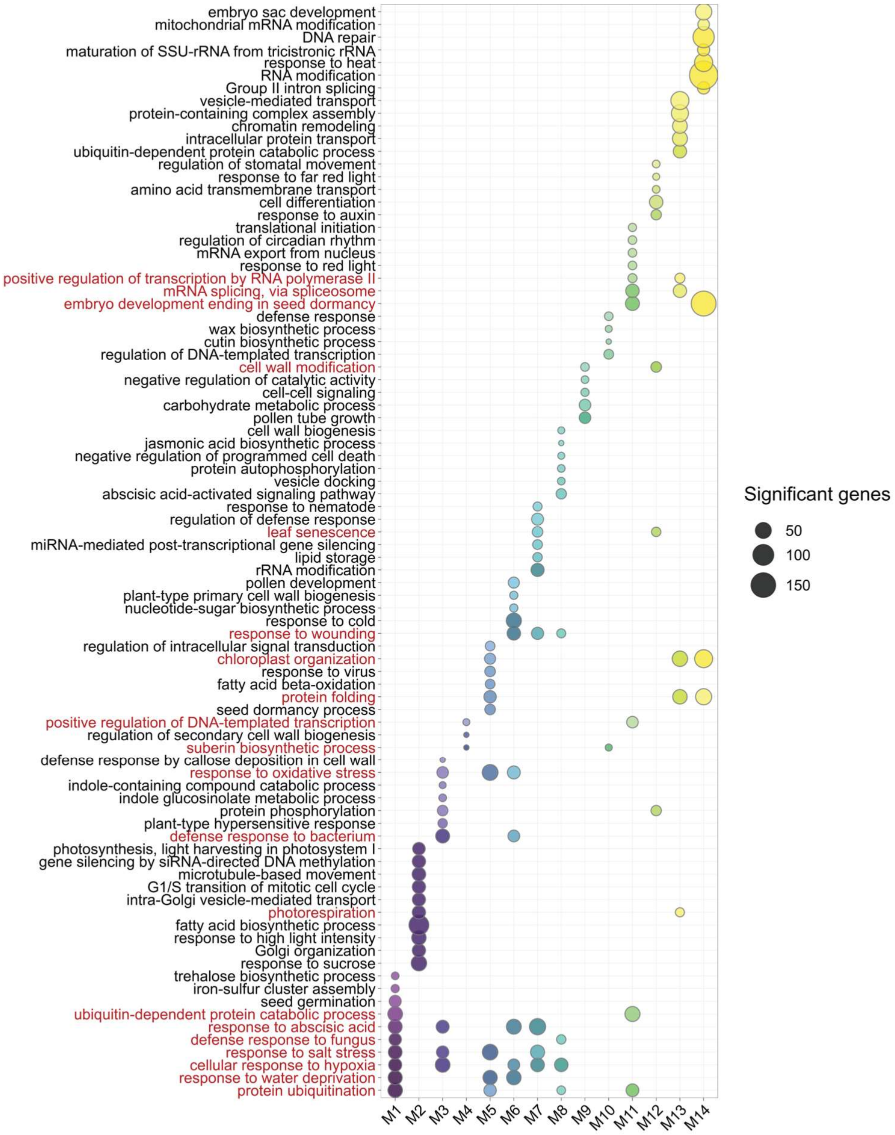


**Fig. S7**. Heatmap showing the top ten GO terms enriched in each gene module. Color intensity indicates -log2 (FDR). Red-colored text indicates GO terms shared across multiple modules.

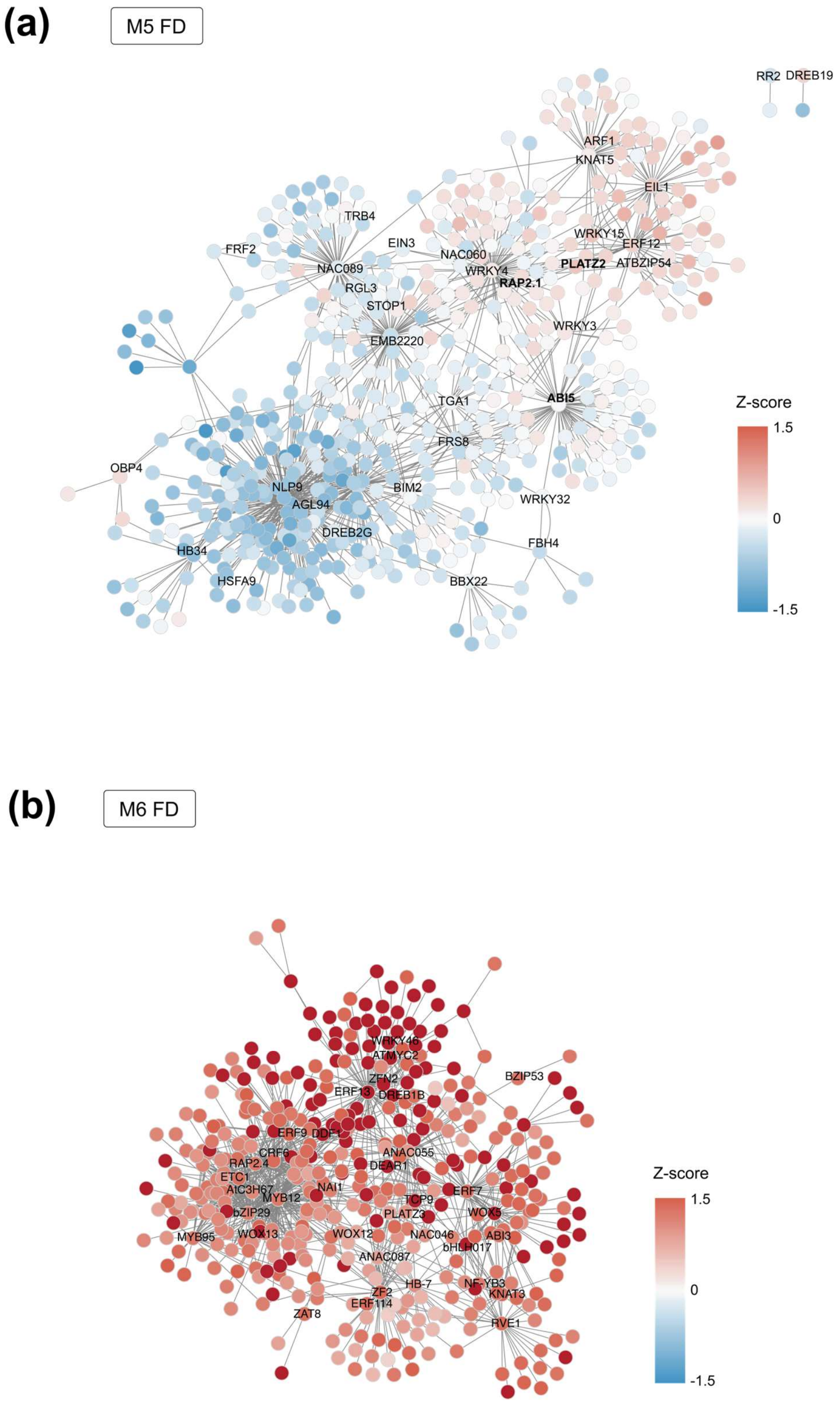


**Fig. S8**. Gene expression dynamics of GRN of M5 and M6 modules in response to FD. **(a)** Z-score changes of genes in the M5 module in response to FD. **(b)** Z-score changes of genes in the M6 module in response to FD. Nodes corresponding to major transcription factors (TFs) are shown in each network.

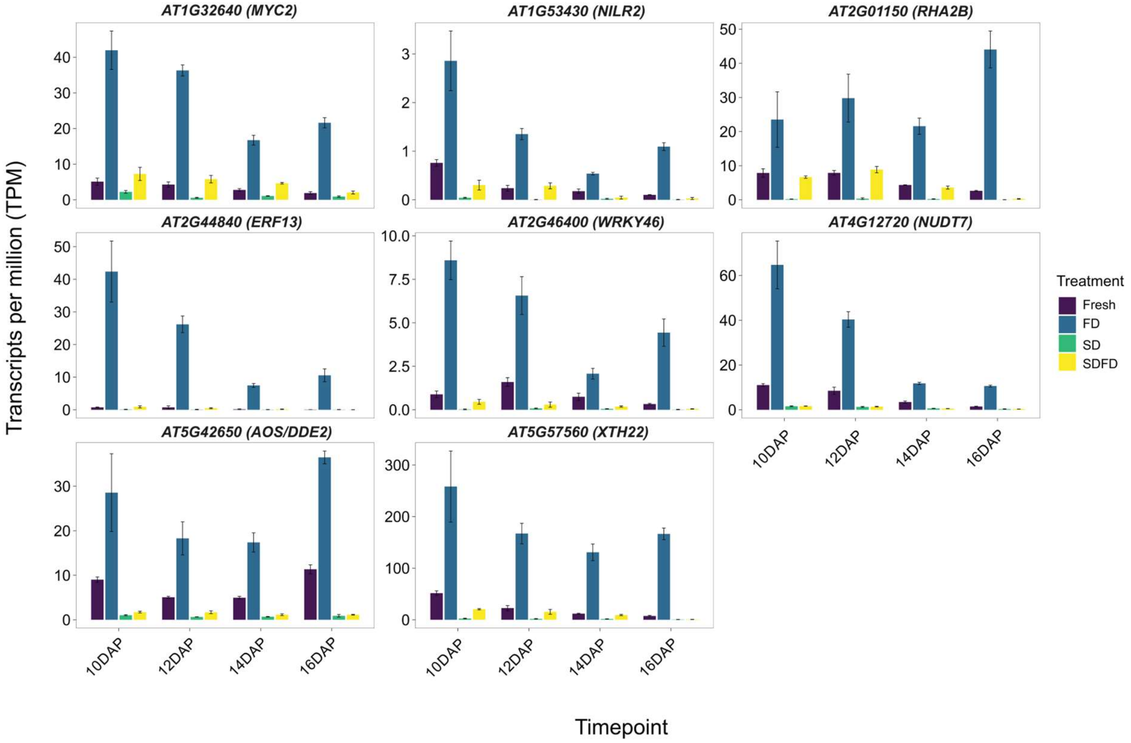


**Fig. S9**. Transcriptional profile of eight genes showing contrasting patterns under FD and SD. Error bars indicate standard error calculated from three replicates (n=3).

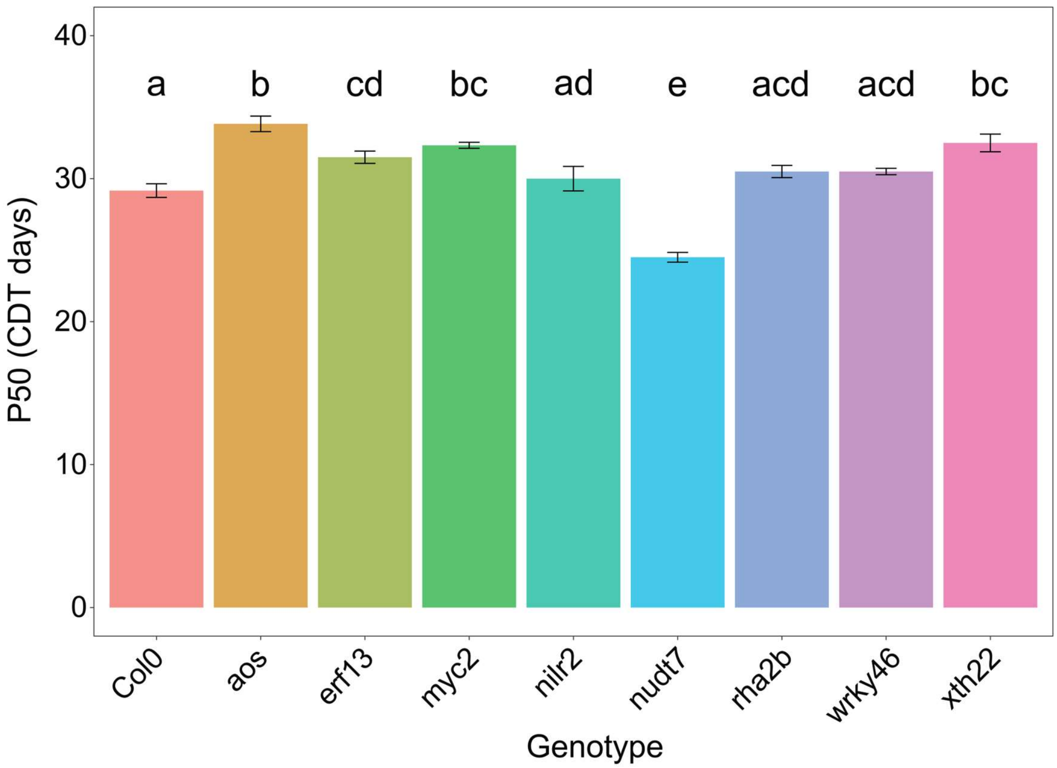


**Fig. S10**. Barplot showing the longevity (p50) of the seeds of different T-DNA lines following a controlled deterioration test (CDT) at 38°C and 75% RH. The mutant genes represent the six genes that overlap between FD up and SD/SDFD down, i.e., genes with contrasting responses to the drying regimes. The wild type (Col-0) was used as a control. Genotype effects on the p50 were determined using the Kruskal-Wallis test followed by Dunn's test with multiple corrections. Significant differences are indicated using letters. Error bars indicate standard error calculated from six replicates (n=6).

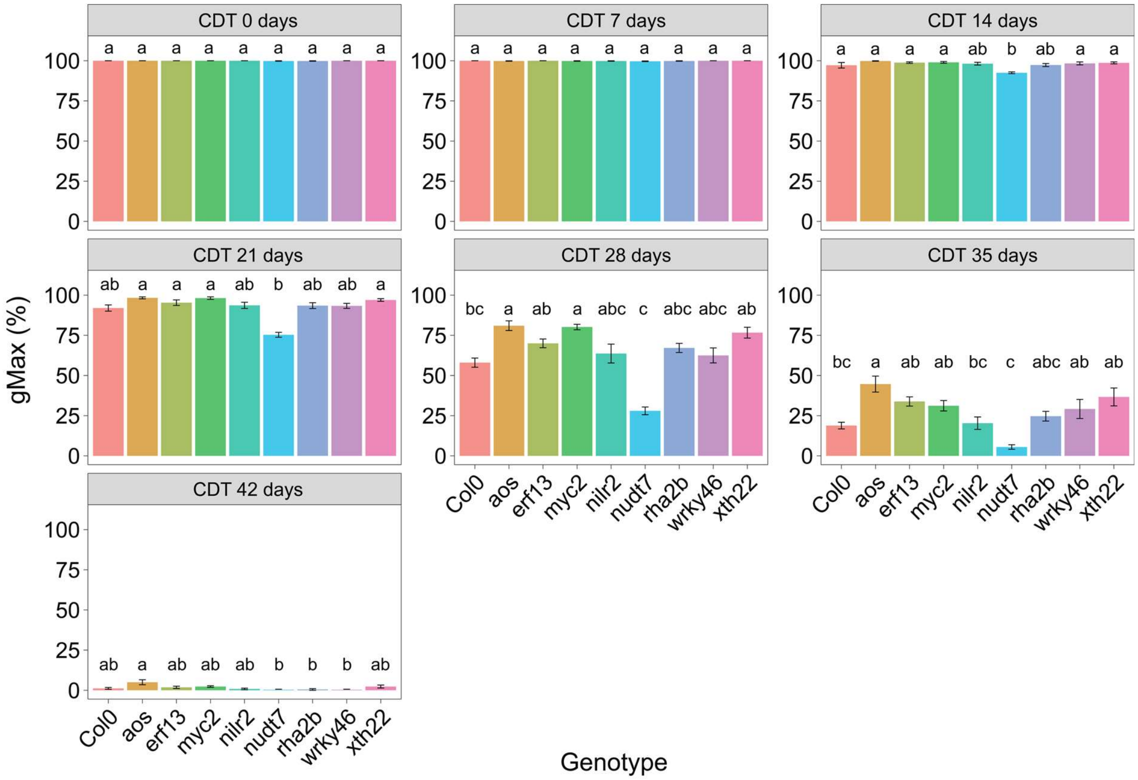


**Fig. S11**. Maximum germination percentage (gMax, %) after 0, 7, 14, 21, 35, and 42 days of controlled deterioration test (CDT) at 38°C and 75% RH. Genotype effects on both gMax were determined using the Kruskal-Wallis test followed by Dunn's test with multiple corrections. Significant differences are indicated using letters. Error bars indicate standard error calculated from six replicates (n=6) over the CDT time points.

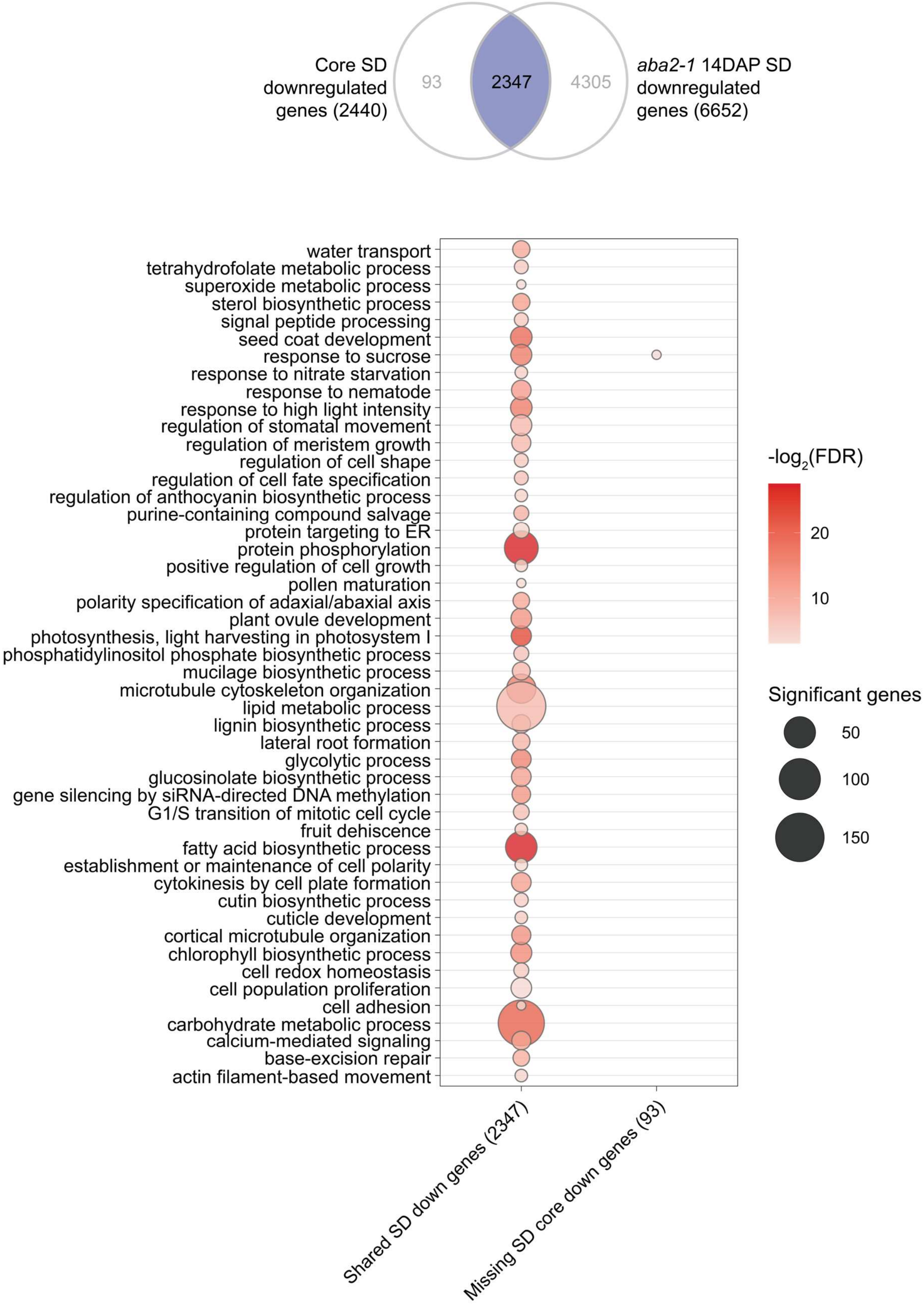


**Fig. S12**. GO enrichment analysis of genes overlapping between Col-0 core SD downregulated and *aba2-1* 14 DAP SD downregulated genes. Size of the bubbles indicate -log2(FDR) values for each GO term.

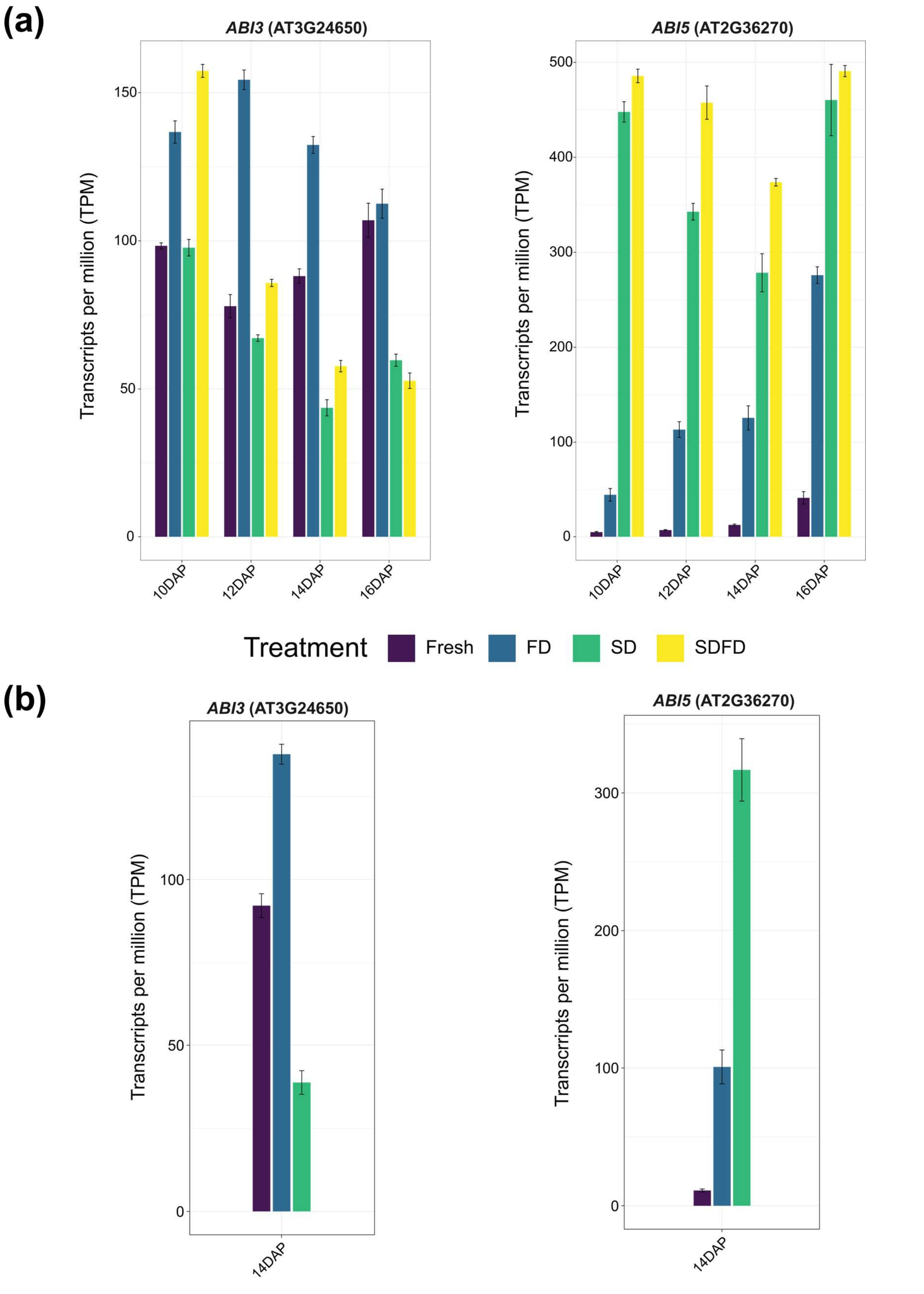


**Fig. S13**. Gene expression level in transcripts per million (TPM) of *ABI3* and *ABI5* genes based on the RNA-seq data. **(a)** Expression of *ABI3* and *ABI5* genes from 10 - 16 DAP in wild type (Col-0). **(b)** Expression of *ABI3* and *ABI5* genes at 14 DAP in the *aba2-1* mutant. Error bars indicate standard error calculated from three replicates (n=3).

**(a)**

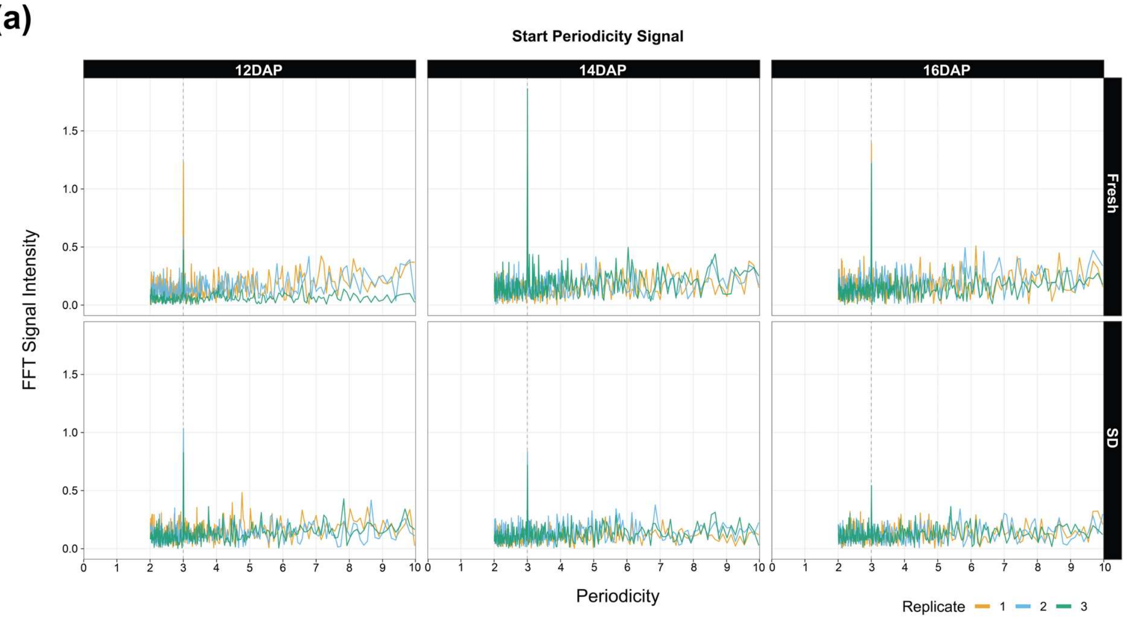


**(b)**

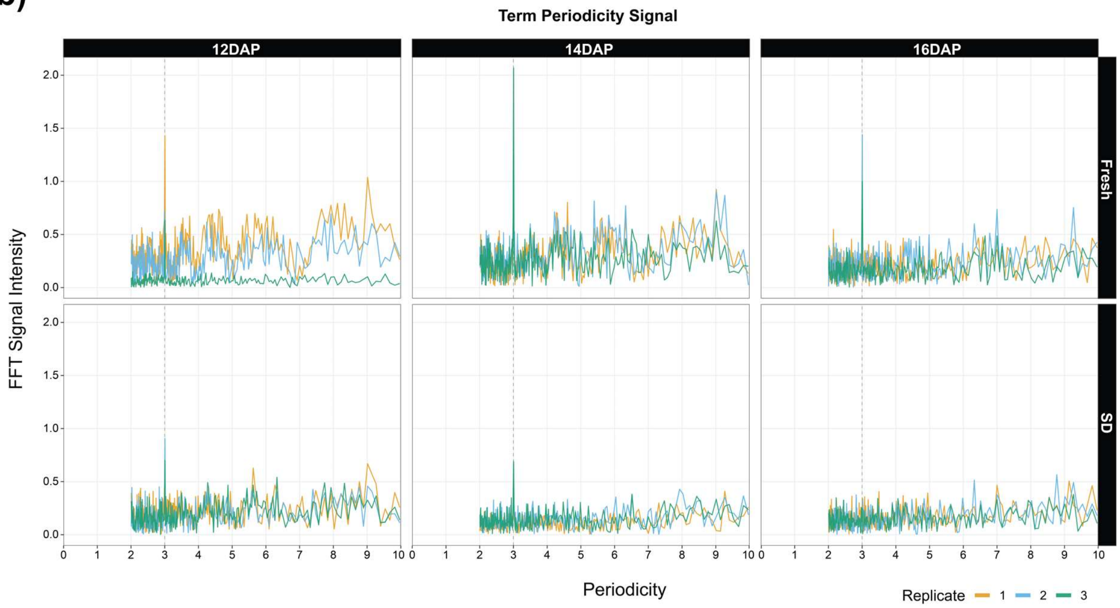


**Fig. S14**. The Fast Fourier transformation (FFT) signal near the **(a)** start and **(b)** stop codon regions. Colors indicate the replicates (n=3).

**(a)**

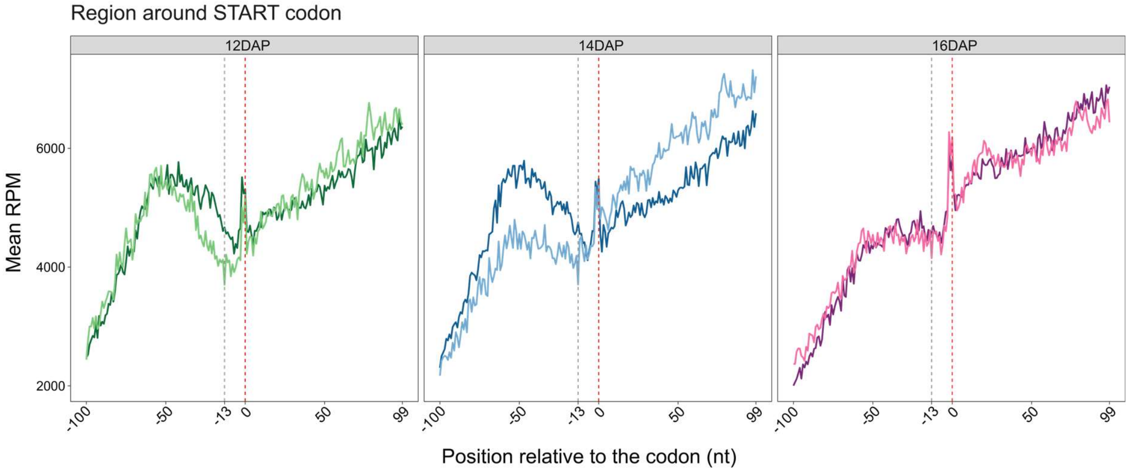


**(b)**

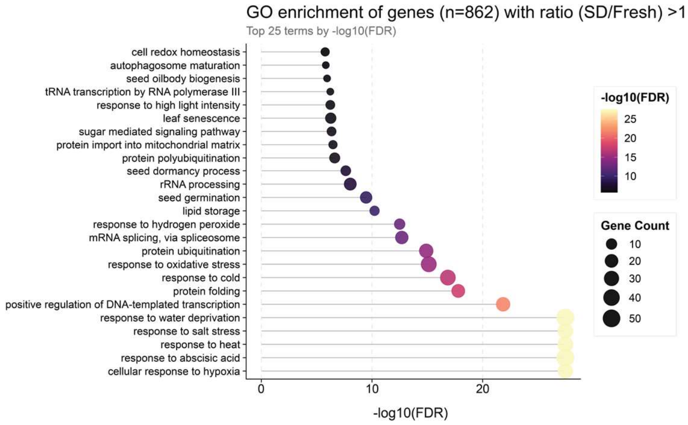


**Fig. S15**. (**a**) Metagene profile near the START codon of genes (n = 862) with a –13 nt read count ratio > 1 between SD and Fresh samples (read count at –13 nt in SD sample/ read count at –13 nt in Fresh) at 12, 14, and 16 DAP. (**b**) Top 25 GO terms enriched in these genes. Color indicates the -log10 (FDR) value, and size indicates the number of significant genes in each GO category.

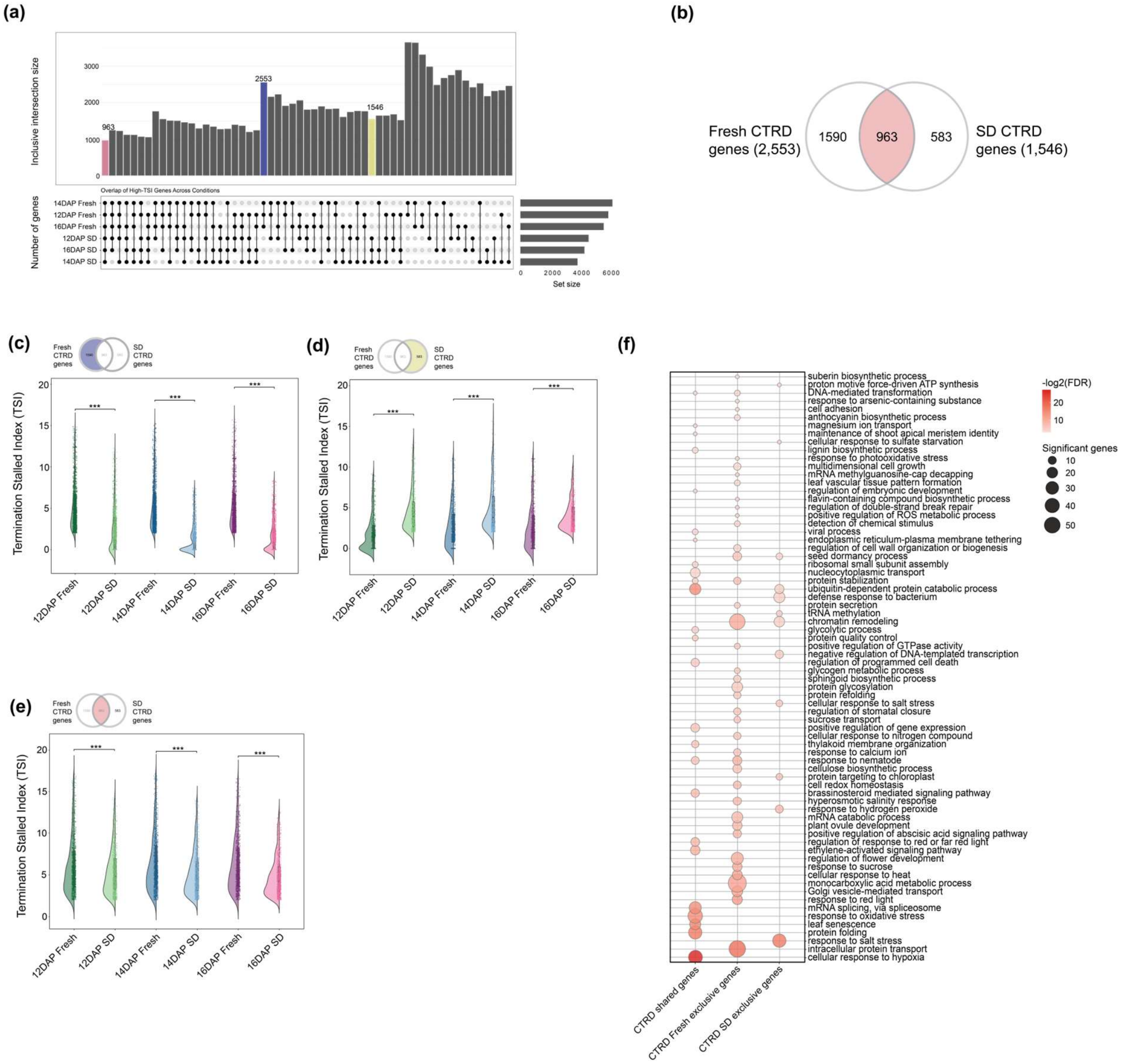


**Fig. S16.** Comparison of genes undergoing CTRD (TSI >= 2) between Fresh and SD samples. (**a**) Upset plot showing the overlap in CTRD genes between different timepoint-treatment combinations. Bars highlighted in red, blue, and yellow indicate CTRD genes shared between all, Fresh, and SD samples, respectively. Gene counts are inclusive. (**b**) Venn diagram showing overlap between genes undergoing CTRD in Fresh and SD samples. (**c**) Distribution of Termination Stalled Index (TSI) of CTRD genes unique to Fresh (left) and SD (right) samples over different timepoint-treatment combinations. (**d**) Distribution of Termination Stalled Index (TSI) of CTRD genes unique to SD. (**e**) Distribution of Termination Stalled Index (TSI) of CTRD genes shared between Fresh and SD. n = 3, biological replicates for all samples except 12 DAP Fresh, where n = 2. Statistical significance was determined using the Wilcoxon rank sum test. (**f**) GO terms enriched in CTRD genes that are specific to Fresh or SD or shared between both.

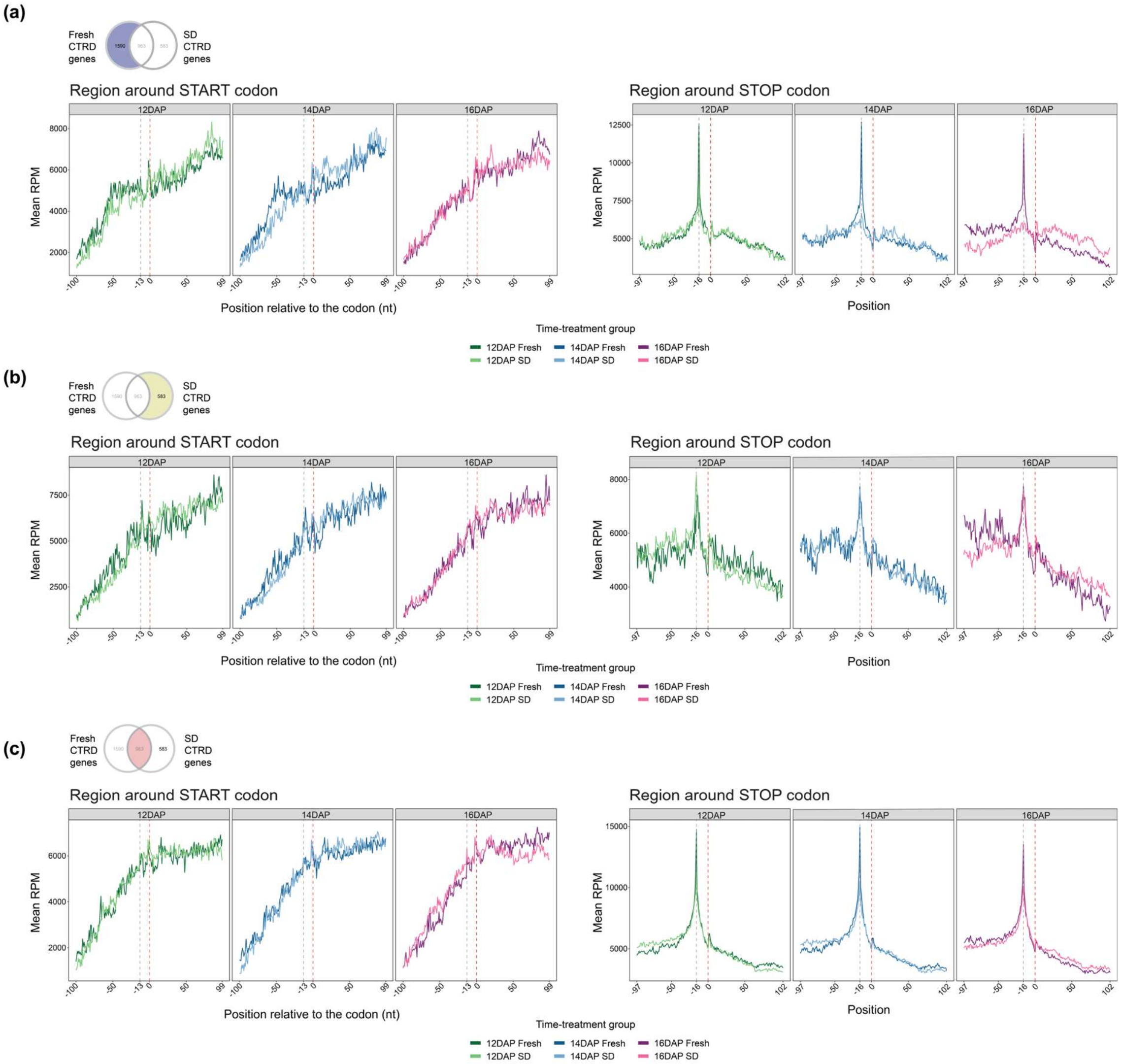


**Fig. S17.** Metagene profiles around start (left) and stop codons (right) of gene undergoing CTRD (TSI>2). (**a**) Metagene profiles of CTRD genes unique to Fresh samples. (**b**) Metagene profiles of CTRD genes unique to SD samples. (**c**) Metagene profiles of CTRD genes shared between Fresh and SD samples.
